# The cIAP ubiquitin ligases sustain type 3 γδ T and innate lymphoid cells during aging to allow normal cutaneous and mucosal responses

**DOI:** 10.1101/2020.05.14.094334

**Authors:** John Rizk, Urs M. Mörbe, Rasmus Agerholm, Isabel Ulmert, Elisa Catafal Tardos, Darshana Kadekar, Maria V. Baglioni, Monica Torrellas Viñals, Vasileios Bekiaris

**Author notes:** Corresponding author: Vasileios Bekiaris.

## Abstract

Environmental and molecular cues early in life are often associated with the permanent shaping of our immune system during adulthood. Although increasing, our knowledge of the signaling pathways that operate in early life and their temporal mode of action is limited. Herein, we demonstrate that the cellular inhibitor of apoptosis proteins 1 and 2 (cIAP1/2), which are E3 ubiquitin ligases and master regulators of the nuclear factor-kappa B (NF-κB) pathway, function during late neonatal and prepubescent life to sustain interleukin(IL)-17-producing gamma delta T cells (γδT17) and group 3 innate lymphoid cells (ILC3). We show that cell-intrinsic deficiency in cIAP1/2 at 3-4 weeks of life leads to downregulation of the transcription factors cMAF and RORγt, and failure to enter cytokine-induced cell cycle. This is followed by progressive loss of γδT17 cells and ILC3 while mice are aging. Mice deficient in cIAP1/2 have severely reduced γδT17 cells and ILC3, present with suboptimal γδT17 responses in the skin, lack small intestinal isolated lymphoid follicles and cannot control intestinal bacterial infection. Mechanistically, these effects appear to be dependent on overt activation of the non-canonical NF-κB pathway. Our data identify the cIAP E3 ubiquitin ligases as critical early life molecular switches for establishing effective type-3 immunity during aging.

## Introduction

The neonatal period is the time when our immune system is imprinted with life-long functional characteristics that maintain immunity to infection and prevent autoimmune pathology. Microbial colonization, and developmentally regulated transcriptional programs cooperate to shape innate and adaptive lymphocytes into distinct specialized lineages that co-exist in equilibrium and respond ad hoc (Eberl, 2016). Failure to convey these environmental and molecular cues during neonatal life, often results in irreversible dysfunction later on. Hence, early dysbiosis impairs type-3 immunity and potentiates susceptibility to type-2 driven allergy (Cahenzli *et al*, 2013) . Similarly, blockade of key signaling pathways during neonatal life can permanently change cellular niches (Kadekar *et al*, 2020). Therefore, elucidating the molecular signatures that operate early in life is of great importance for understanding how immunity develops.

Mouse γδ T cells present a well-established example of an immune population that is heavily dependent on an unperturbed neonatal period. In this regard, intestinal intraepithelial (IE) γδ T cells develop during neonatal and prepubescent life through butyrophylin-driven interactions with the epithelia (Di Marco Barros *et al*, 2016) . This provides a necessary defense mechanism against infection within the IEL compartment (Hoytema van Konijnenburg *et al*, 2017). Lamina propria (LP) interleukin(IL)-17-producing γδ T cells establish mixed type-3 and type-1 transcriptional programs within the first week of life through the transcription factor STAT5 (Kadekar *et al*, 2020) . Thus, early life establishment of the γδT17 compartment is critical to protect from neonatal and adult infections (Chen *et al*, 2020; Sheridan *et al*, 2013). In a similar manner, impaired microbial colonization of the ocular or oral mucosa results in drastically altered IL-17-producing γδ T (γδT17) cell numbers in the conjunctiva (St. Leger *et al*, 2017) and cervical lymph nodes (LN) (Fleming *et al*, 2017). Again, paucity in such γδT17 cell populations is associated with impaired anti-microbial responses in eye and oral cavity (Conti *et al*, 2014; St. Leger *et al*, 2017), and resistance to pathogenic inflammation (Cai *et al*, 2011; Sandrock *et al*, 2018; McGinley *et al*, 2020).

The innate lymphoid cell (ILC) compartment is also dependent on early life events, while their function during the neonatal period is critical for the establishment of the intestinal immune system (Spits *et al*, 2013). In this regard, although dysbiosis does not affect ILC development, it results in altered transcriptional and epigenetic profiles of all ILC subsets (Gury-BenAri *et al*, 2016). Similar to LP γδT17 cells, group 3 ILC (ILC3) acquire expression of the transcription factor Tbet and the type-1 cytokine interferon-γ (IFN-γ) during neonatal life, which allows them to clear intracellular bacterial infections (Klose *et al*, 2013) . Moreover, group 2 ILC (ILC2) undergo an IL-33-dependent maturation step in the neonatal lung, allowing their cytokine responsiveness in adult mice (Steer *et al*, 2020). Importantly, ILC3 induce the maturation of intestinal cryptopathces into isolated lymphoid follicles (ILFs) during the first 3-4 weeks of life (Kiss *et al*, 2011; Kruglov *et al*, 2013). Evidently, perturbations of γδT17 and ILC3 development during the early stages of life will have a substantial impact on the quality of immunity while aging. The molecular pathways that control the transition of these cells from neonatal life to adolescence and adulthood are poorly understood.

The E3 ubiquitin ligases cellular inhibitor of apoptosis protein (cIAP)1 and 2 (cIAP1/2) catalyze both degradative lysine(K)-48 and stabilizing K-63 ubiquitination and act as the main molecular switches for the activation of the canonical and non-canonical nuclear factor-kappa B (NF-κB) pathway (Silke & Meier, 2013) . The presence of cIAP1/2 downstream of TNF receptor 1 (TNFR1) determines whether a cell will initiate the canonical NF-κB pathway or die by apoptosis or necroptosis in response to TNF (Annibaldi & Meier, 2018). They achieve this by ubiquitinating receptor interacting kinase-1 (RIPK1) (Silke & Meier, 2013) . However, cIAP1/2 are mostly recognized as negative regulators of the non-canonical NF-κB pathway. Hence, in all cell types cIAP1/2 associate in a heterocomplex with TNF receptor associated factor (TRAF)2, TRAF3 and NF-κB-inducing kinase (NIK), whereby they induce K-48 ubiquitination of NIK, resulting in its continuous proteasomal degradation (Zarnegar *et al*, 2008; Varfolomeev *et al*, 2007; Vince *et al*, 2007). Breakdown of the TRAF2-TRAF3-cIAP1/2-NIK complex either following ligation of TNF superfamily receptors that recruit TRAF2-TRAF3 in their intracellular domain (e.g. TNFR2, LTβR, CD40) or by cIAP1/2 depletion, liberates NIK, which initiates the cascade necessary for nuclear translocation of the non-canonical NF-κB transcription factors RelB and p52 (Vallabhapurapu *et al*, 2008; Matsuzawa *et al*, 2008)

In the present study we demonstrate a necessary role for cIAP1/2 in sustaining γδT17 cells and ILC3 at the late neonatal and prepubescent stages of life, and thus impacting the magnitude of inflammatory and anti-bacterial immune responses. Deficiency in cIAP1/2 begun to have an impact only during late neonatal life by reducing expression of the lineage defining transcription factors cMAF and RORγt, which was followed by an apparent block in cytokine-induced proliferation. When animals entered prepubescence and early adolescence, cIAP1/2 deficiency resulted in progressive loss γδT17 cells. This was independent of TNFR1 induced canonical NF-κB or cell death. In contrast, cIAP1/2-deficient prepubescent γδT17 cells displayed enhanced nuclear translocation of RelB, which demonstrates evidence of overt activation of the non-canonical NF-κB pathway. Intestinal ILC3 also relied on intact cIAP1/2 during the same time period, with their numbers being drastically reduced in adulthood. Paucity in ILC3 coincided with ILF involution. Mice with targeted deletion of cIAP1/2 in γδT17 cells and ILC3 responded sub-optimally to cutaneous inflammatory challenge and failed to control intestinal bacterial infection.

## Results

### Paucity of γδT17 cells in the absence of the E3 ubiquitin ligases cIAP1 and cIAP2

Using acute, SMAC mimetic (SM) driven antagonization and in vitro techniques, we showed before that cIAP1/2 are important for T_H_17 differentiation through modulation of the non-canonical NF-κB pathway (Rizk *et al*, 2019). In order to understand the in vivo importance of cIAP1 and cIAP2 in RORγt-expressing immune cells, we crossed *Rorc*-Cre (RORγt^CRE^) mice (Eberl & Litman, 2004) with mice that were floxed for *Birc2* (cIAP1^F/F^) and knocked out for *Birc3* (cIAP2^-/-^) (Gardam *et al*, 2011). This generated mice with RORγt-driven deletion of cIAP1 (referred to thereafter as ΔIAP1) and generalized deletion of cIAP2 (referred to thereafter as ΔIAP2), as well as the corresponding Cre-negative littermate controls (wild-type; WT) (Fig 1A). ΔIAP1 and ΔIAP1/2 mice were viable, produced offspring at expected rates, and did not develop any observable spontaneous disease phenotypes. They contained a full set of lymph nodes (inguinal, brachial, axillary and mesenteric) and Peyer’s patches indicating unperturbed lymphoid tissue development, while total numbers of CD4^+^ T and B cells were normal but γδ T were slightly elevated (Fig S1A).

**Figure 1.**
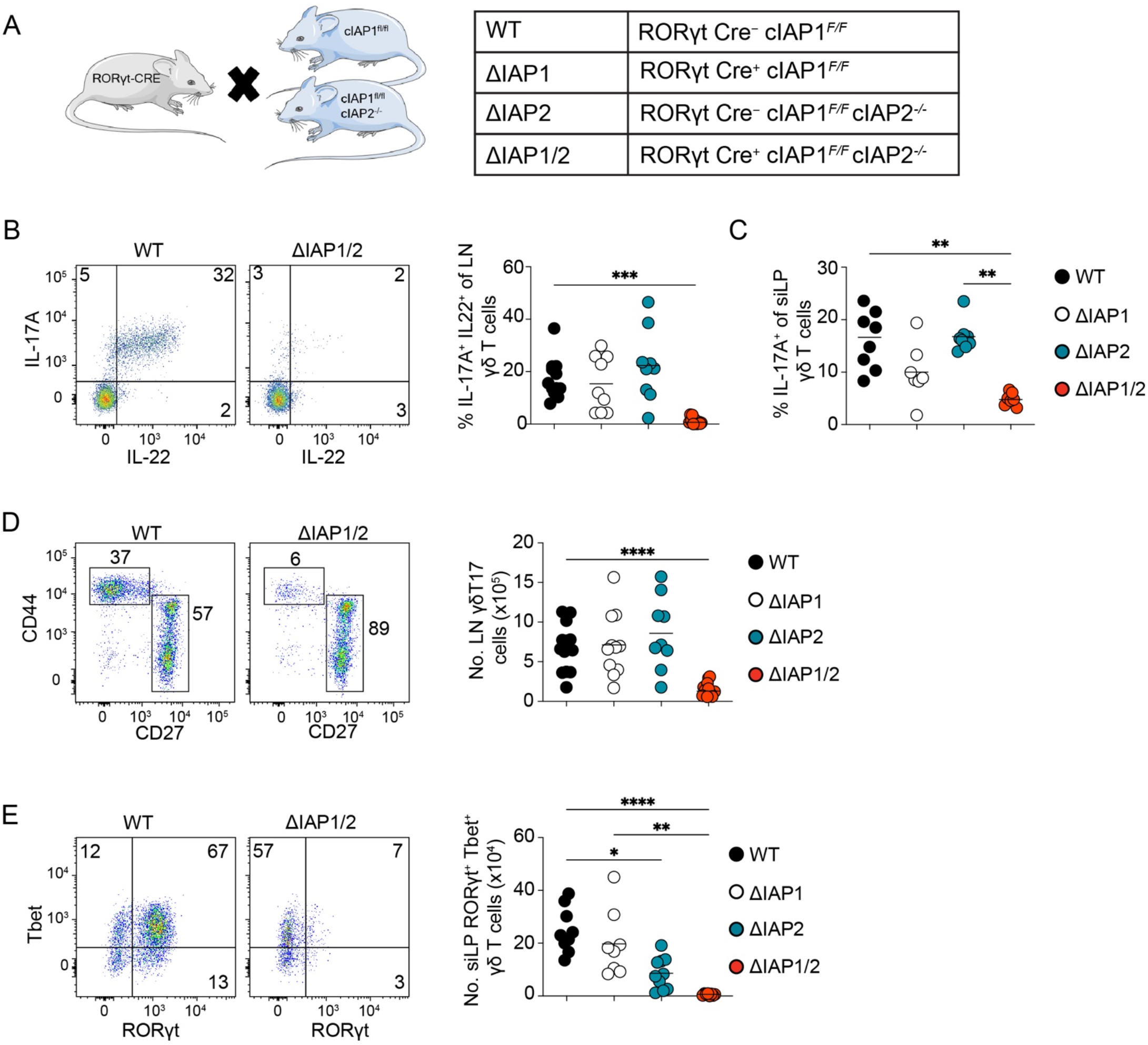
cIAP1 and cIAP2 are required for the homeostasis of γδT17 cells in the LNs and intestinal lamina propria. (A) Graphical representation of the different mouse strains generated by crossing RORγt^CRE^ mice to cIAP1^F/F^ or cIAP1^F/F^ cIAP2^-/-^ mice. Representative flow cytometric analysis (B) and frequency (B-C) of IL-17^+^ IL-22^+^ cells within γδ T cells in the LNs (B) or (C) IL-17^+^ cells within γδ T in the siLP. (D) Representative flow cytometric analysis (dot plots) and numbers (graph) of γδT17 cells in the LNs of adult WT, ΔIAP1, ΔIAP2 and ΔIAP1/2 mice. (E) Representative flow cytometric analysis (dot plots) and numbers (graph) of RORγt^+^ Tbet^+^ γδ T cells in the siLP of WT, ΔIAP1, ΔIAP2 and ΔIAP1/2 mice. In graphs, each symbol represents a mouse, and lines represent the mean, data is pool of 4 experiments in (B) or 5 experiments in (C-E). *P < 0.05, **P < 0.01, ***P < 0.001, ****P < 0.0001 using Kruskal-Wallis test with Dunn’s correction.

We next analyzed some of the major IL-17-producing populations in lymph node (LN) and small intestinal and colonic lamina propria (siLP; cLP). Compared to littermate controls, ΔIAP1/2 mice produced slightly elevated levels of IL-17A within the CD4^+^TCRβ^+^ compartment in the LN (pool of inguinal, brachial, axillary) but not the gut (Fig S1B), suggesting that in these animals, steady-state production of IL-17A by CD4^+^ T cells is not defective. Staining for IL-22 following overnight stimulation with IL-23 yielded the same answer (Fig S1C). However, there was a marked reduction in γδ-associated IL-17A and IL-22 production in LN (Fig 1B) and IL-17A in the gut (Fig 1C). This was accompanied by a dramatic loss in LN TCRγδ^+^CD27^−^CD44^hi^CCR6^+^, which are the IL-17-producing γδ T cells (Ribot *et al*, 2009; Haas *et al*, 2009) (Fig 1D and Fig S1D), and gut Tbet^+^RORγt^+^ (Kadekar *et al*, 2020) (Fig 1E) γδT17 cell numbers. Although cIAP1 and cIAP2 individually did not contribute to this phenotype in the LN, ΔIAP2 mice had significantly reduced Tbet^+^RORγt^+^ γδ T cell numbers in the gut (Fig 1E). We additionally found significantly reduced γδT17 cells in the lungs of ΔIAP2 mice (Fig S1E). Similar to γδT17 cells, there were significantly reduced non-CD4 IL-17-producing lymphocytes in the LNs of ΔIAP1/2 mice (Fig S1F).

In the skin, CD3^lo^Vγ5^−^TCRγδ^+^CCR6^+^ cells, which represent the γδT17 population (Haas *et al*, 2009, 2012), were also reduced significantly in the absence of cIAP1 and cIAP2 (Fig 2A). When we analyzed the two major γδT17 subpopulations (Vγ4^+^ versus Vγ4^−^; Vγ nomenclature by Heilig and Tonegawa; (Heilig & Tonegawa, 1986)), we found that in the skin cIAP1 but not cIAP2 was required for Vγ4^−^ cells, whereas the Vγ4-expressing population was only affected by the absence of both cIAP1 and cIAP2 (Fig 2B). In the LN, we did not observe differential regulation of either Vγ4^+^ or Vγ4^−^ cells (Fig S2A). Collectively, this data suggests that cIAP1/2 are important for the development and/or homeostatic maintenance of γδT17 cells. Our findings additionally pinpoint a differential and non-redundant role of cIAP1 and cIAP2 in these cells that is organ and subset specific. In this regard, whereas skin γδT17 cells depended more on cIAP1, gut γδT17 cells depended more on cIAP2.

**Figure 2.**
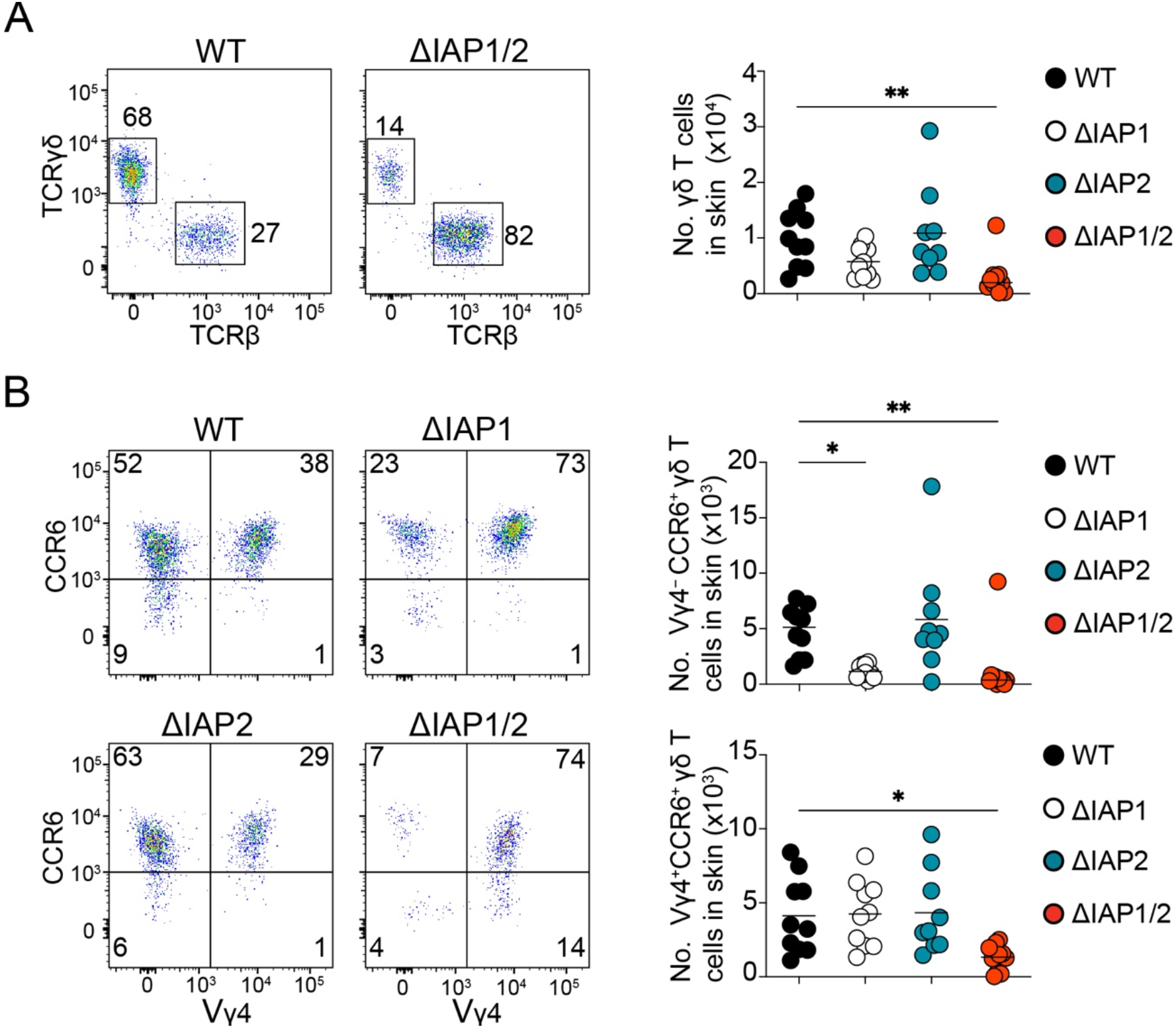
cIAP1 and cIAP2 are non-redundantly required for the homeostatic maintenance of γδT17 cell subsets in the skin. Representative flow cytometric analysis (dot plots) and numbers (graphs) of (A) total γδ T cells or (B) Vγ4^+^ and Vγ4^-^ CCR6^+^ γδ T cells in the skin of WT, ΔIAP1, ΔIAP2 and ΔIAP1/2 mice. In graphs, each symbol represents a mouse, and lines represent the mean, data is pool of 5 experiments in (A-B). *P < 0.05, **P < 0.01 using Kruskal-Wallis test with Dunn’s correction.

### Cell-intrinsic requirement for cIAP1 and cIAP2 in γδT17 cells

Next, we investigated whether the defect we observed in ΔIAP1/2 mice was cell-intrinsic. To this end we set up mixed bone marrow (BM) chimeras where WT CD45.1^+^CD45.2^+^ hosts were sub-lethally irradiated and reconstituted with a mixture of 1:1 CD45.1^+^ WT and CD45.2^+^ ΔIAP1/2 BM cells (Fig 3A). We found that, under these conditions, WT LN γδT17 cells outcompeted their ΔIAP1/2 counterparts (Fig 3B), indicating the phenotype we observed in intact mice was cell intrinsic. Interestingly, CD27^+^ γδ T cells derived from ΔIAP1/2 BM were slightly less competitive than WT (Fig 3B). In contrast, both CD3^−^ populations and B cells from ΔIAP1/2 BM were more competitive than their WT counterparts (Fig 3C). This indicated that the reduced competitiveness of γδT17 and CD27^+^ γδ T cells was not due to defective ΔIAP1/2 BM reconstitution. We could not reconstitute gut RORγt^+^Tbet^+^ γδT17 cells irrespective of the BM source (Fig 3D), suggesting that this population requires either thymus-originated γδ T cells or a neonatal microenvironment to develop fully. In contrast, lack of cIAP1 and cIAP2 did not impinge on the reconstitution of gut Tbet^+^RORγt^−^ γδ T cells (Fig 3E). Likewise, we could only recover WT LN γδT17 cells when we reconstituted ΔIAP1/2 hosts with WT or a 1:1 mix of WT and ΔIAP1/2 BM (Fig S3A-B), while CD27^+^ γδ T cells from WT or ΔIAP1/2 BM cells were equally competitive (Fig S3B). As before, we could not reconstitute RORγt^+^Tbet^+^ γδT17 cells in the gut (Fig S3C).

**Figure 3.**
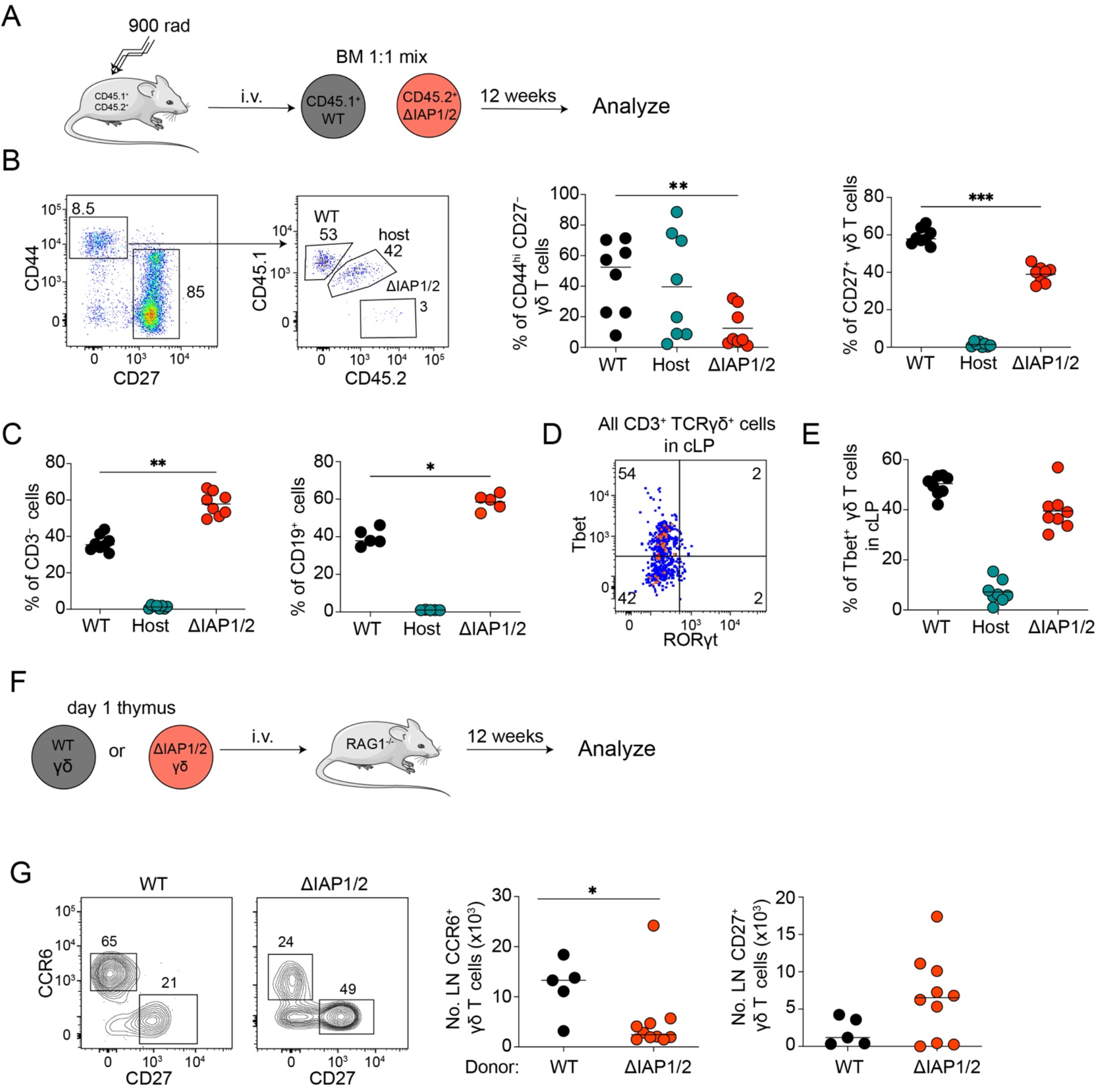
cIAP1 and cIAP2 are intrinsically required for the homeostasis of γδT17 cells. (A) Graphical representation of the experimental setup for competitive bone marrow experiments. (B) Representative flow cytometric analysis (dot plots) and frequency (graphs) of WT (CD45.1^+^), host (CD45.1^+^ CD45.2^+^) or ΔIAP1/2 (CD45.2^+^) -derived γδT17 and CD27^+^ γδ T cells within γδ T cells population in the LNs of reconstituted hosts. (C) Frequency of WT (CD45.1^+^), host (CD45.1^+^ CD45.2^+^) or ΔIAP1/2 (CD45.2^+^) - derived CD3^-^ and CD19^+^ cells in the LNs of reconstituted hosts. (D) Flow cytometric analysis and (E) frequency of RORγt^+^ and Tbet^+^ γδ T cells in the cLP of host mice follwing bone marrow reconstitution. (B-E) In graphs, each symbol represents a mouse, and lines represent the mean, data is pool of 3 experiments. *P < 0.05, **P < 0.01, ***P < 0.01 using one-way ANOVA with Tukey’s correction. (F) Graphical representation of the experimental setup for transfer of neonatal γδ T cells to RAG1^-/-^ recipients. (G) Flow cytometric analysis (contour plots) and numbers (graphs) of CCR6^+^ CD27^-^ or CD27^+^ γδ T cells in the LNs of RAG1^-/-^ hosts after transfer of neonatal γδ T cells from WT or ΔIAP1/2 pups. In graphs, each symbol represents a mouse, and lines represent the mean, data is pool of 3 experiments (G). *P < 0.05 using Mann-Whitney test.

As γδT17 cells develop perinatally in the thymus and undergo a rapid neonatal re-programming within the tissues they localize at, we reasoned that if generated from BM stem cells, they might have different developmental or homeostatic requirements for cIAP1 and cIAP2. To address this issue, we purified γδ T cells from the thymi of 1-day old WT or ΔIAP1/2 mice and transferred them to RAG1^−/−^ recipients (Fig 3F). We found that 12 weeks post transfer, the γδT17 cell compartment was reconstituted in the LN, however, we recovered significantly more WT than ΔIAP1/2 cells (Fig 3G). As with the BM chimeras, we could not reconstitute intestinal RORγt^+^Tbet^+^ γδT17 cells, suggesting that this population requires a neonatal microenvironment (Fig S3D). Reconstitution of CD27^+^ γδ T cells was independent of cIAP1 and cIAP2 (Fig 3G). Taken together our data show that γδT17 cells require cIAP1 and cIAP2 intrinsically.

### The impact of cIAP1 and cIAP2 on γδT17 cells is independent of TNF induced canonical NF-κB and cell death

In addition to preventing spontaneous activation of the non-canonical NF-κB pathway, cIAP1/2 are necessary to convey the canonical NF-κB downstream of TNFR1 whereas in their absence, TNF-TNFR1 interactions can lead to RIPK1-mediated cell death via apoptosis or necroptosis (Annibaldi & Meier, 2018). Mice deficient in TNFR1 had an intact γδT17 cell population (Fig S4A), suggesting that the canonical NF-κB pathway downstream of TNFR1 is not responsible for the phenotype of ΔIAP1/2 mice. Since TNF is highly upregulated during the weaning reaction (Al Nabhani *et al*, 2019), we next investigated whether TNF induced cell death played a role. To achieve this, we initially analyzed mice that were deficient in cIAP2 and expressed a ubiquitin-associated (UBA) domain mutant form of cIAP1 unable to K48 ubiquitylate and suppress RIPK1 (Annibaldi *et al*, 2018). Thus, these mice are more sensitive to TNF induced cell death (Annibaldi *et al*, 2018). In UBA-mutant mice, γδT17 cells were not affected (Fig S4B), suggesting that these cells are not susceptible to death by homeostatic levels of TNF. In order to test this directly in ΔIAP1/2 mice, we began injecting 1-week old neonates with neutralizing anti-TNF antibody and until animals were 12-week old (Fig S4C). We could not rescue the γδT17 population in either gut or LNs (Fig S4D-E), indicating that TNF induced death is unlikely to play a major role in regulating these cells in the absence of cIAP1/2. Therefore, TNF-TNFR1 interactions are not responsible for the ΔIAP1/2 phenotype, suggesting that overt activation of the non-canonical NF-κB pathway could play a key role.

### cIAP1 and cIAP2 are required for γδT17 cell cycle progression and expression of cMAF and RORγt during aging

In order to assess the impact of cIAP1/2 on embryonic γδT17 cell development, we enumerated thymic cell numbers in newborn ΔIAP1/2 mice and found them similar to littermate controls (Fig 4A), despite efficient deletion of *Birc2* (cIAP1) (Fig S5A). Production of IL-17A/F were unchanged at this stage (Fig S5B). This suggested that the major impact of cIAP1/2 occurs post-embryonically. We therefore tracked LN γδT17 cells, defined phenotypically as CD27^−^CD44^hi^, during neonatal, post-neonatal (average weaning time at 3 weeks) and adult life (mating age of 8 weeks). We did not find any differences in cell numbers until week 5 of age (Fig 4B). This suggested that cIAP1 and cIAP2 are only required to sustain γδT17 numbers following weaning. After week 5, ΔIAP1/2 γδT17 cells failed to expand and began to decline progressively during aging (Fig 4B). In order to confirm that the cells are missing from adult life and have not converted to a non-γδT17 population, we crossed ΔIAP1/2 with the ROSA26-LSL-RFP strain, so that RFP permanently marks all current and “ex” RORγt-expressing cells. We found no evidence of γδT17 conversion to other populations (Fig 4C). This data suggested that cIAP1/2 regulate a checkpoint in early adult life that manifests during aging.

**Figure 4.**
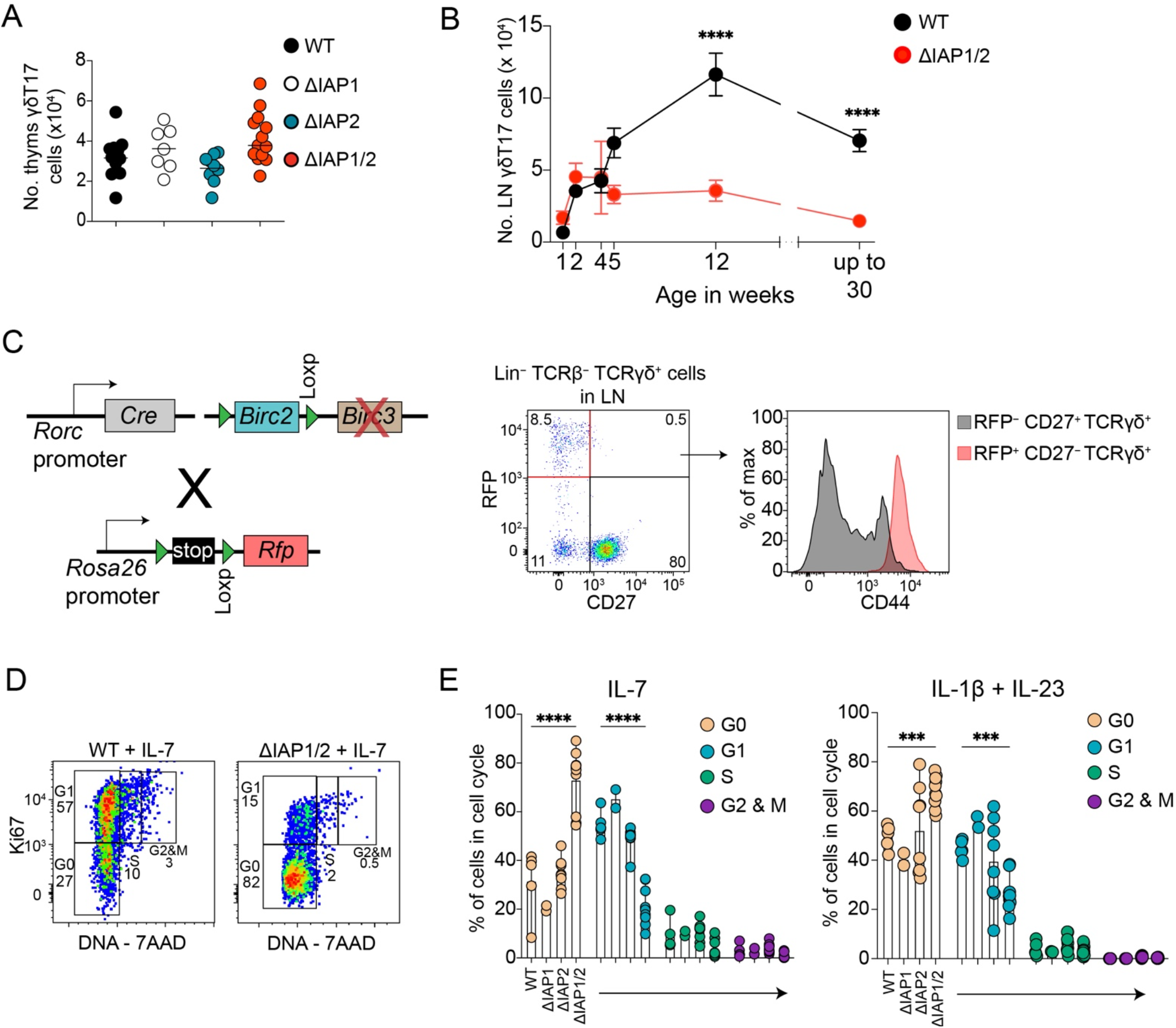
cIAP1 and 2 double deficient γδT17 cells are lost progressively during aging and fail to progress through cell cycle. (A) Numbers of γδT17 cells in the thymi of 1-day old WT, ΔIAP1, ΔIAP2 and ΔIAP1/2 pups. In graph, each symbol represents a mouse, and lines represent the mean, data is a pool of 3 experiments. (B) Numbers of γδT17 cells in the LN of WT and ΔIAP1/2 at the indicated timepoints. Each symbol represents the mean amalgamated data from each timepoint and the error bars represents the SEM. ****P < 0.01 using Two-way ANOVA with Holm-Sidak correction. (C) Graphical representation of the genetic makeup of the ΔIAP1/2 mice when crossed to the ROSA26-LSL-RFP strain, and representative flow cytometry analysis of LN γδT17 cells from ΔIAP1/2 x ROSA26-LSL-RFP mice. (D) Representative flow cytometric analysis and (E) frequency of cells in G0, G1, S or G2/M cell cycle stages within γδT17 cells that were ex-vivo cultured with the indicated cytokines for 48 hours. In graphs, each symbol represents a mouse, and bars represent the mean, data is pool of 3 experiments. ***P < 0.001, ****P < 0.0001 using two-way ANOVA with Tukey’s correction.

We next investigated what this checkpoint was. The inability of the cells to increase in numbers during aging, raised the hypothesis that cIAP1/2 may be regulating responsiveness to cytokines that induce proliferation. We thus isolated 4-week old LN cells and treated them in vitro with IL-7 or a combination of IL-1β+IL-23. We found that ΔIAP1/2 γδT17 cells were slower in entering cell cycle with most cells stuck in G0 (Fig 4D-E). This defect in proliferation was also observed following stimulation with either IL-2 or TCR cross-linking (Fig S5C). Therefore, cIAP1/2 are important for γδT17 cell cycle progression.

We have shown before that ablation of cIAP1/2 in T cells downregulates cMAF, a lineage determining transcription factor for γδT17 cells (Zuberbuehler *et al*, 2019), in a NIK- and RelB-dependent mechanism (Rizk *et al*, 2019). We thus hypothesized that lack of cIAP1/2 may influence expression of cMAF. In newborn thymus expression of cMAF as well as RORγt was unchanged (Fig S6A). At week 1 after birth we observed a slight reduction in the expression of RORγt and cMAF (Fig S6B). However, at week 3 of age, expression of RORγt and cMAF was significantly reduced (Fig 5A). Furthermore, we observed a modest but significant reduction in IL-17A production by 3-week old ΔIAP1/2 γδT17 cells (Fig 5B). In the intestine, RORγt^+^ γδ T cells express high levels of CD127 (IL-7Rα) and intermediate levels of CD45 (Fig 5C). Due to lack of other reliable surface markers to identify these cells in the gut, we gated TCRγδ^+^CD45^int^CD127^+^ cells and quantified numbers as well as expression of RORγt and cMAF. Similar to the LNs, numbers in the siLP did not change in 4-week old ΔIAP1/2 mice (Fig 5D), however there was a significant reduction in the levels of RORγt and cMAF (Fig 5E).

**Figure 5.**
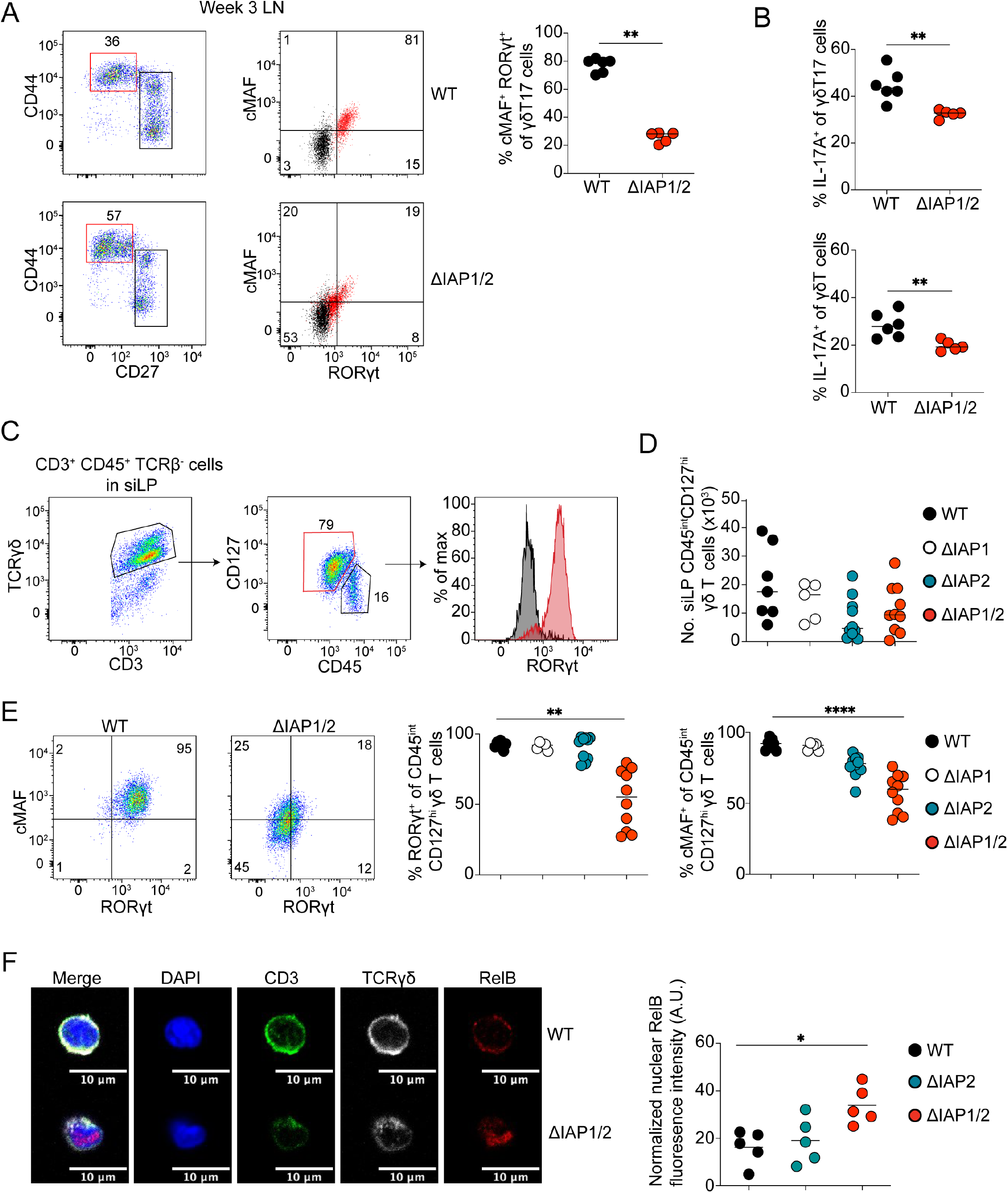
cIAP1 and 2 maintain the transcription factor landscape of γδT17 cells in the lymph nodes and intestinal lamina propria. (A) Flow cytometric analysis (dot plots) and frequency (graphs) of RORγt^+^ cMAF^+^ cells within γδT17 cells from the LNs of 3-week-old WT and ΔIAP1/2 mice. (B) frequency of IL-17^+^ cells within γδT17 cells (top) or within all γδ T cells (bottom) from the LNs of 3-week-old WT and ΔIAP1/2 mice. In graphs, each symbol represents a mouse, and the line represent the mean, data is pool of 2 experiments. **P < 0.01 using Mann-Whitney test. (C) Flow cytometric analysis showing the gating strategy and expression of RORγt by CD45^int^ CD127^+^ γδ T cells in siLP of adult WT mice. (D) Numbers of CD45^int^ CD127^+^ γδ T cells in the siLP of 4-week-old WT, ΔIAP1, ΔIAP2 and ΔIAP1/2 mice. (E) Flow cytometric analysis (dot plots) and quantification (graphs) of RORγt or cMAF expression by CD45^int^ CD127^+^ γδ T cells in the siLP of 4-week-old WT, ΔIAP1, ΔIAP2 and ΔIAP1/2 mice. In graphs, each symbol represents a mouse, and the line represent the mean, data is pool of 4 experiments. (F) Representative immunofluorescent microscopy analysis of γδT17 cells (CD27^-^ TCRγδ^+^ cells) and normalized nuclear RelB fluorescence intensity in sorted from the LNs of 4-week-old WT or ΔIAP1/2. Images are representative of four independent experiments. *P < 0.05, **P < 0.01, ****P < 0.001 using using Kruskal-Wallis test with Dunn’s correction.

Furthermore, we investigated the expression of RelB in newborn thymic γδ T cells and found a significant upregulation of *Relb* mRNA in ΔIAP1/2 γδT17 but not in CD27^+^ γδ T cells (Fig S6C). We additionally examined the extent of RelB nuclear translocation in γδT17 cells from 4-week old WT, ΔIAP2 and ΔIAP1/2 mice. We found that the levels of nuclear RelB in ΔIAP1/2 γδT17 cells were significantly higher by comparison to WT cells (Fig 5F), arguing for a role of the non-canonical NF-κB pathway in downregulation of RORγt and cMAF. Therefore, cIAP1/2 are required during late neonatal life in order to maintain expression of the transcription factors RORγt and cMAF and to sustain normal γδT17 numbers.

### Inflammation partially restores γδT17 responses in the absence of cIAP1 and cIAP2

Next, we investigated whether cytokines that activate γδT17 cells could regulate expression of RORγt and cMAF from 4-week old mice. Culture with IL-7 did not influence expression of either transcription factors (Fig 6A-C), however, a combination of IL-1β and IL-23 resulted in partial restoration of RORγt but not cMAF in ΔIAP1/2 γδT17 cells (Fig 6A-C). Interestingly, IL-1β+IL-23 resulted in downregulation of cMAF in WT cells (Fig 6A-B). We additionally observed that the ΔIAP1/2 γδT17 cells that acquired RORγt, were the cells that entered G1 in response to IL-1β+IL-23 and to a lesser extent in response to IL-7 (Fig S7). In the imiquimod(IMQ)-driven psoriasiform dermatitis model, IL-23, IL-1β and IL-7 drive γδT17 cell expansion as well as production of IL-17 and IL-22 in the LN and skin (Michel *et al*, 2012; Cai *et al*, 2011, 2014). We thus treated WT, ΔIAP1, ΔIAP2, and ΔIAP1/2 4-week old mice with IMQ for seven days and assessed expression of RORγt and cMAF in γδT17 cells. We found that IMQ treatment partially restored expression of both RORγt and cMAF in ΔIAP1/2 γδT17 cells in the LNs (Fig 6D), suggesting that inflammation can rescue the ΔIAP1/2 phenotype.

**Figure 6.**
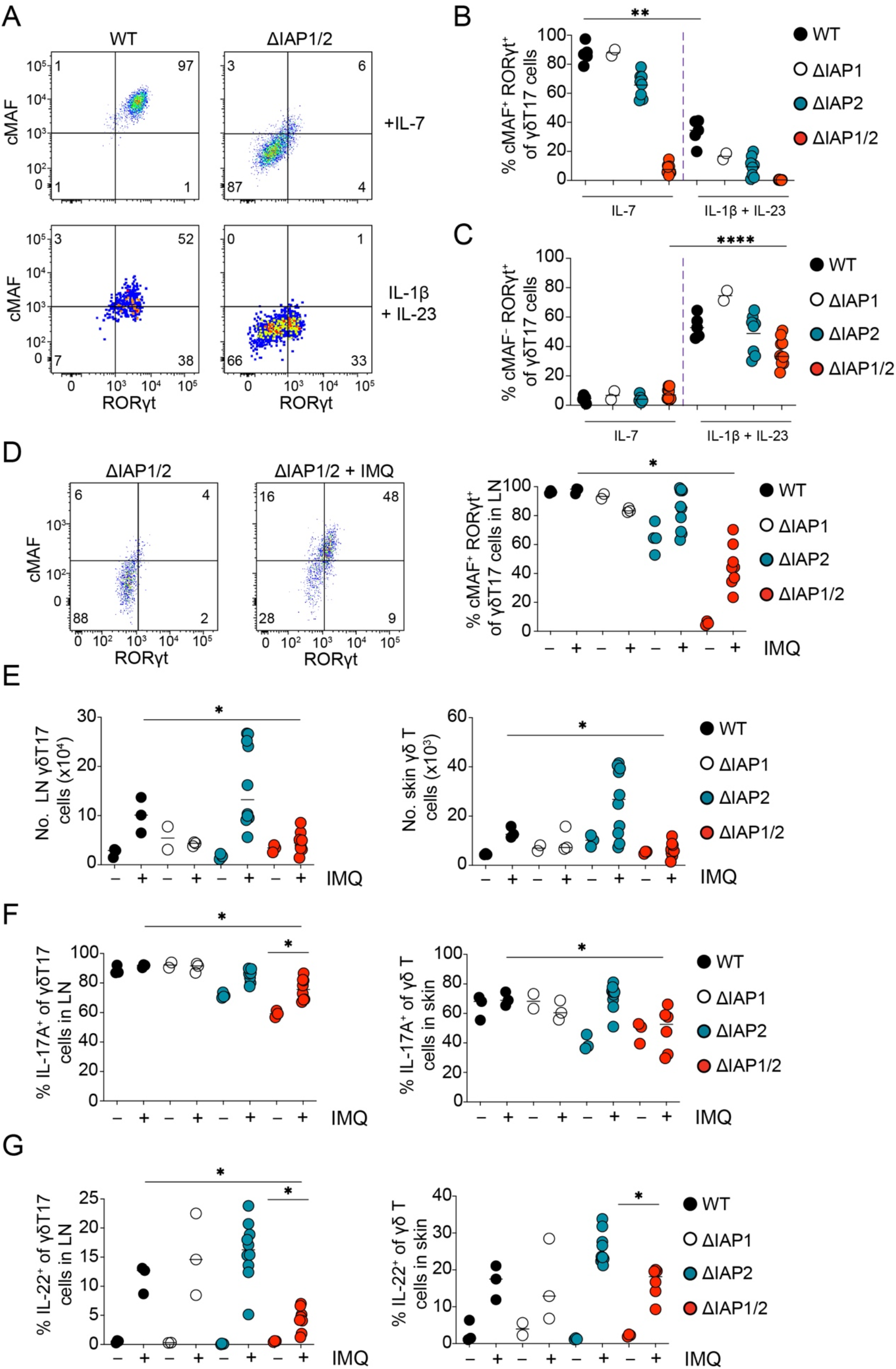
Inflammation partially overcomes cIAP1 and cIAP2 deficiency in γδT17 cells. (A) Representative flow cytometric analysis and (B-C) quantification of RORγt and cMAF expression by γδT17 cells from the LNs of 4-week-old WT or ΔIAP1/2 mice following ex-vivo culture with the indicated cytokines for 48 hours. In graphs, each symbol represents a mouse, and the line represent the mean, data is pool of 3 experiments. (D) Representative flow cytometric analysis (dot plot) and quantification (graph) of RORγt and cMAF expression by γδT17 in LNs of 4-week-old control or IMQ-treated ΔIAP1/2 mice. (E) Numbers of γδT17 cells in the LNs (right) or skin (left) of 4-week-old control or IMQ-treated WT, ΔIAP1, ΔIAP2 or ΔIAP1/2 mice. (F-G) Frequency of IL-17A^+^ (F) or IL-22^+^ (G) cells within γδT17 cells in the LNs or skin of 4-week-old control or IMQ-treated WT, ΔIAP1, ΔIAP2 or ΔIAP1/2 mice. In graphs, each symbol represents a mouse, and the line represent the median, data is pool of 3 experiments. *P< 0.05, **P < 0.01, ****P < 0.0001 using Mann-Whitney test.

We then investigated whether rescue of RORγt and cMAF was sufficient for ΔIAP1/2 γδT17 cells to mount an immune response. We observed that despite an increase in Ki67 expression (Fig S8A), ΔIAP1/2 γδT17 numbers did not increase in either the LNs or skin (Fig 6E). Evaluation of cytokine production revealed substantial regional differences between LN and skin in ΔIAP1/2 mice. Thus, whereas in the LN, ΔIAP1/2 γδT17 cells increased (albeit significantly less than their WT, ΔIAP1 and ΔIAP2 counterparts) their production of IL-17A following IMQ treatment, this was not the case in the skin (Fig 6F). In contrast, IL-22 production in LNs was significantly reduced while it was relatively normal in the skin (Fig 6G). The CD4^+^ T cell response to IMQ was not defective and slightly stronger in ΔIAP1/2 mice (Fig S8B). The extent of skin inflammation, as measured by epidermal thickening, was not different between ΔIAP1/2 and control mice, reflecting both the partial rescue of the γδT17 as well as the slightly exaggerated CD4^+^ T cell response (Fig S8C).

The data suggest that although at a young age cIAP1/2 regulate proliferation, transcriptional stability and cytokine production, strong inflammatory stimuli can, to a certain extent, overcome this regulatory checkpoint and revive γδT17 cell responses. These results additionally indicate that the extrathymic expression and biological impact thereafter of RORγt and cMAF can be dynamic and under the control of multiple microenvironment cues.

### cIAP1 and cIAP2 are required for intestinal ILC3 during aging and for sustaining ILF integrity

ILC3 share many functional characteristics and transcription factor requirements with γδT17 cells, including constitutive expression of RORγt and cMAF (Zuberbuehler *et al*, 2019; Parker *et al*, 2019). We therefore investigated the impact of cIAP1 and cIAP2 deficiency in intestinal ILC3 populations. Similar to γδT17 cells, LP Tbet^+^ and Tbet^−^ ILC3 numbers were reduced in ΔIAP1/2 mice (Fig 7A-C and Fig S9A). As expected ILC2 numbers were not affected (Fig S9B). Next, we tested whether ΔIAP1/2 ILC3 converted to an RORγt^−^ population and hence performed a lineage tracing experiment using the RORγt-RFP reporter mice described above. We found that within the ILC population, there was a 10-fold reduction in RFP^+^ cell numbers, suggesting that in the absence of cIAP1 and cIAP2, there is loss of ILC3 rather than conversion to a non-ILC3 population (Fig S9C). Similar to γδT17 cells, we found that ILC3 numbers did not expand post weaning (Fig 7D). Next, we investigated whether the ILC3 defect in ΔIAP1/2 mice was cell-intrinsic. To this end, using mixed BM chimeras, we found that WT ILC3 outcompeted their ΔIAP1/2 counterparts (Fig 7E-F), indicating that the phenotype we observed in intact mice was cell-intrinsic. There was equal reconstitution capacity of GATA3-expressing ILC2 derived from WT or ΔIAP1/2 BM (Fig S9D), demonstrating the specificity of the defect within RORγt-expressing populations. We obtained similar results when we reconstituted ΔIAP1/2 hosts with a 1:1 mix of WT and ΔIAP1/2 BM (Fig 7G and Fig S9E). Further, cIAP1 and 2 deficient ILC3 cells were not rescued by treatment with anti-TNF (Fig S9F).

**Figure 7.**
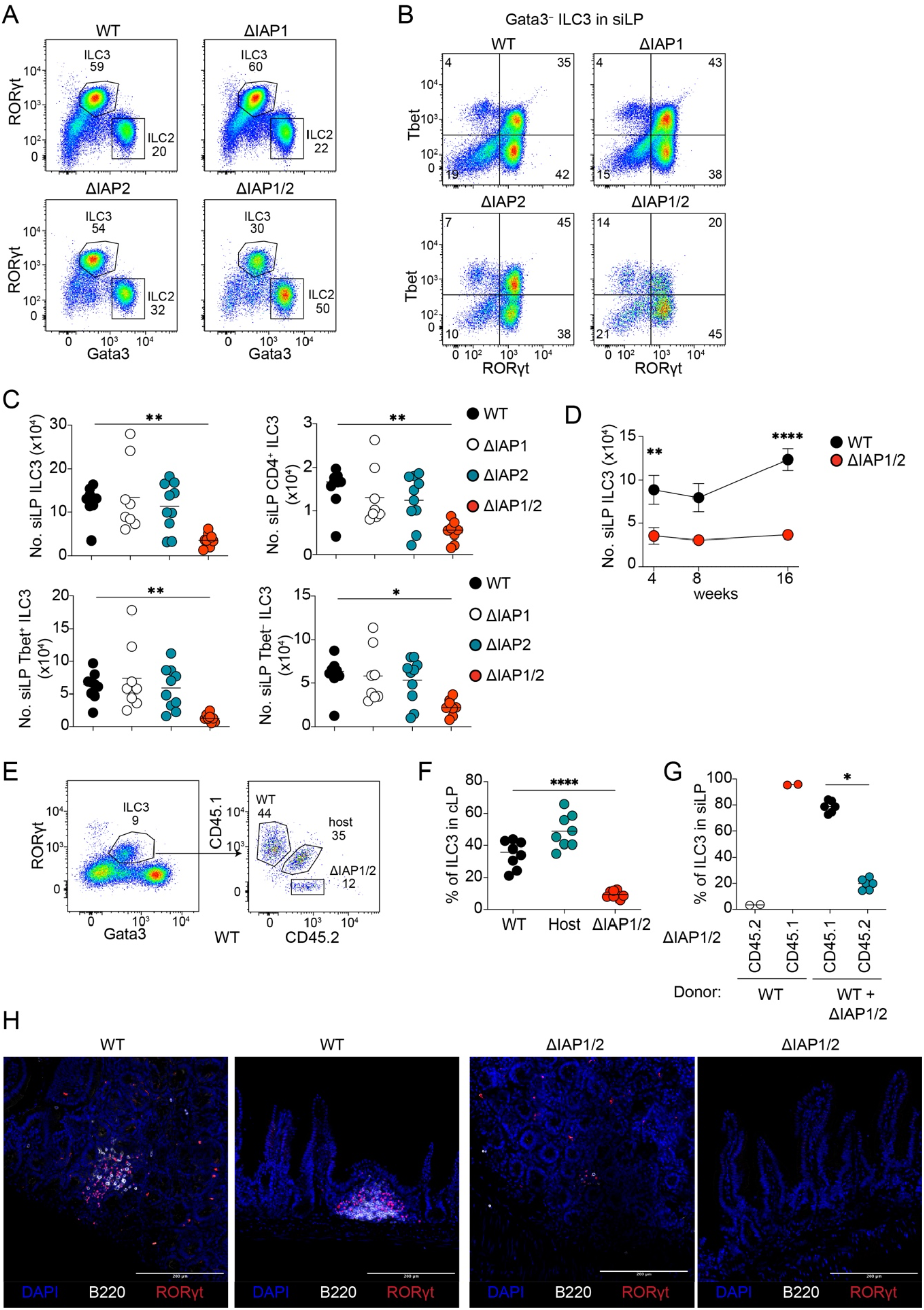
cIAP1 and cIAP2 are intrinsically required for the maintanance of intestinal ILC3 and for ILF integrity. Representative flow cytometric analysis of (A) total CD45^+^ CD3^-^ CD19^-^ CD127^+^ ILCs or (B) CD45^+^ CD3^-^ CD19^-^ CD127^+^ GATA3^-^ cells in the siLP of adult WT, ΔIAP1, ΔIAP2 or ΔIAP1/2 mice. (C) Numbers of total ILC3s, CD4^+^, Tbet^+^ or Tbet^-^ ILC3s in the siLP of adult WT, ΔIAP1, ΔIAP2 or ΔIAP1/2 mice. In graphs, each symbol represents a mouse, and the line represent the mean, data is pool of 5 experiments. *P< 0.05, **P < 0.01 using Kruskal-Wallis test with Dunn’s correction. (D) Numbers of ILC3s in the siLP of WT and ΔIAP1/2 at the indicated timepoints. Each symbol represents the mean amalgamated data from each timepoint and the error bars represents the SEM. **P < 0.01, ****P < 0.0001 using Two-way ANOVA with Holm-Sidak correction. (E) representative flow cytometric analysis (dot plots) and (F) frequency (graph) of WT (CD45.1^+^), host (CD45.1^+^ CD45.2^+^) or ΔIAP1/2 (CD45.2^+^)-derived ILC3 in the cLP of reconstituted hosts. In the graph, each symbol represents a mouse, and lines represent the mean, data is pool of 3 experiments. ***P < 0.01 using one-way ANOVA with Tukey’s correction. (G) Frequency of WT (CD45.1^+^) or ΔIAP1/2(CD45.2^+^)-derived ILC3 in the siLP of bone marrow reconstituted ΔIAP1/2 hosts. In graph, each symbol represents a mouse, and lines represent the mean, data is pool of 2 experiments. *P < 0.05 using Wilcoxon-rank t-test. (H) representative immunofluorescent microscopy images showing ILF structures in distal ileum sections from adult WT or ΔIAP1/2 mice. Images are representative of two independent experiments.

ILC3 are necessary for the maturation of cryptopatches to ILFs during the first weeks of life. ΔIAP1/2 mice had severely defective ILFs (Fig 7H). ILFs in these mice were either absent or reduced in size (Fig 7H). Despite the lack of ILFs, production of IgA was not defective (Fig S9G). Collectively, this data shows that cIAP1 and cIAP2 are necessary for intestinal ILC3 to expand during the post-weaning period, and to induce formation of ILFs.

### cIAP1 and cIAP2 are necessary to protect against *Citrobacter rodentium* infection

It has been demonstrated that ILC3 are important to control infection by the attaching and effacing bacterium *Citrobacter rodentium* (Bauché *et al*, 2020; Guo *et al*, 2015, 2014), a widely used model for human enteropathogenic *E. coli* infections (Silberger *et al*, 2017) .We therefore reasoned that ΔIAP1/2 mice may be defective in mounting a protective response to *C. rodentium*. We infected ΔIAP1/2 mice and their respective controls with 2x10^9^ CFU of *C. rodentium* through oral gavage and followed weight loss as a surrogate marker for disease. We found that by 11 days after infection ΔIAP1/2 mice lost approximately 20% of their body weight, while all other strains did not (Fig 8A). At this time point and due to ethical constraints, all animals were sacrificed and we analyzed bacterial loads and the immune response in the colon. ΔIAP1/2 mice had significantly higher colonic bacterial load than controls (Fig 8B). This was associated with compromised IL-22 production from the ILC3 compartment (Fig 8C-D).

**Figure 8.**
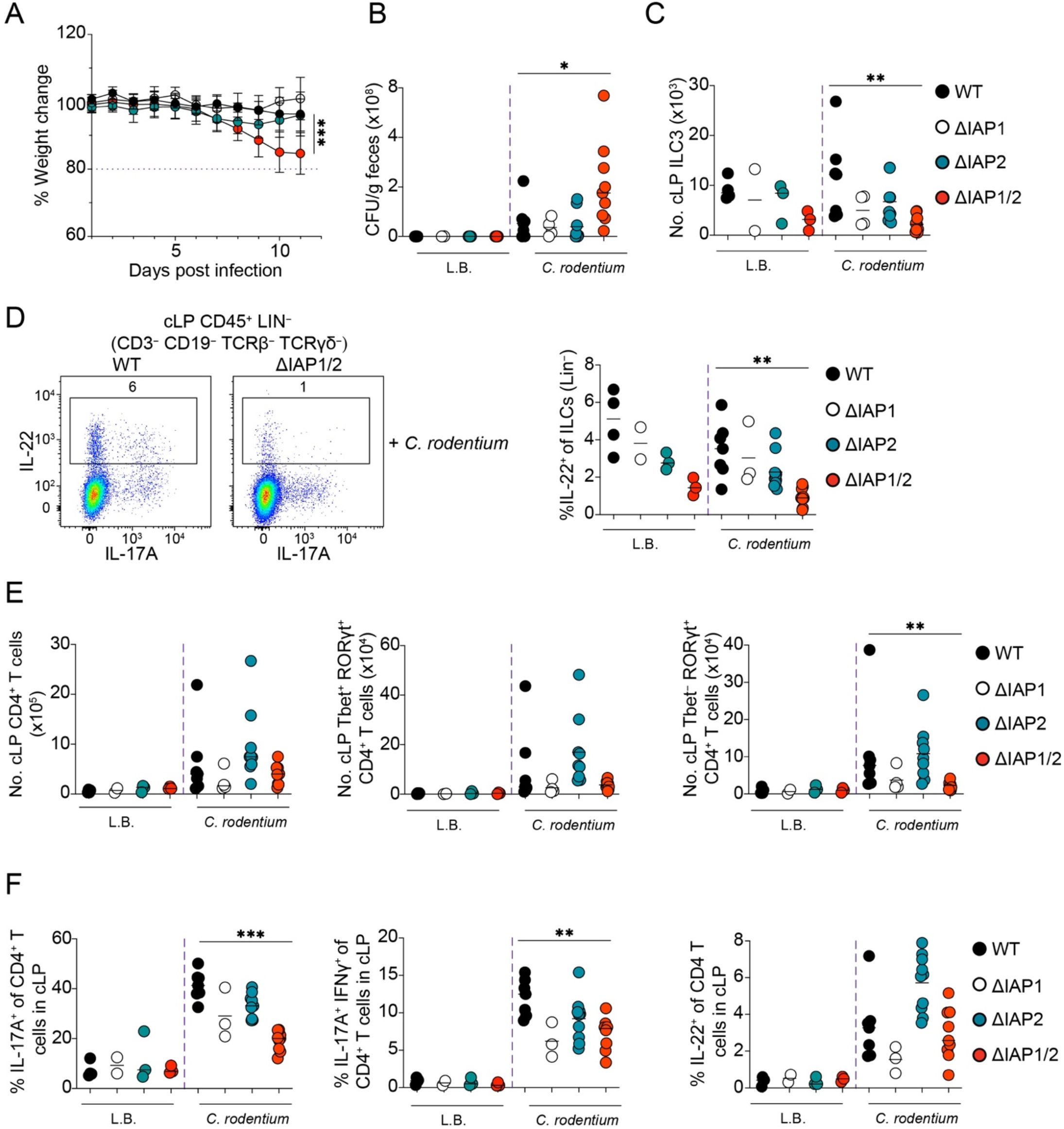
RORγt^Cre+^ cIAP1^F/F^ cIAP2^-/-^ succumb to *Citrobacter rodentium* infections. (A) Percentage body weight change of *C. rodentium* infected WT, ΔIAP1, ΔIAP2 and ΔIAP1/2 mice. Each symbol represents the mean amalgamated data from each timepoint and the error bars represents the SD. ***P < 0.001 using Two-way ANOVA with Holm-Sidak correction. (B) Colony forming units (CFU) of *C. rodentium* in fecal matter of infected and uninfected WT, ΔIAP1, ΔIAP2 and ΔIAP1/2 mice. (C) Numbers of ILC3s in the cLP of infected and uninfected mice. (D) representative flow cytometric analysis (dot blots) and frequency of IL-22^+^ cells within ILCs in the cLP of *C. rodentium* infected WT and ΔIAP1/2 mice. (E) Numbers of total CD4^+^ T cells, Tbet^+^ RORγt^+^ CD4^+^ T cells, and RORγt^+^ Tbet-CD4^+^ T cells in the cLP of infected and uninfected mice. (F) Frequency of IL-17A^+^, IL-17A^+^ IFNγ^+^ or IL-22^+^ cells within CD4^+^ T cells in the cLP of infected and uninfected mice. In graphs, each symbol represents a mouse, and lines represent the median, data is pool of 3 experiments. *P < 0.05, **P < 0.01, ***P < 0.001 using Mann-Whittney test.

Although ILC3 are important to protect from *C. rodentium* infection, a T_H_17 and T_H_22 response is also required, as evidenced by susceptibility of RAG1^−/−^ mice to this pathogen (Silberger *et al*, 2017). We therefore additionally analyzed the CD4^+^ T cell response in the colon. Numbers of total CD4^+^ T cells were not changed in infected ΔIAP1/2 mice (Fig 8E). However, there was a significant reduction in RORγt^+^Tbet^−^ CD4^+^ T cells (Fig 8E), which was accompanied by reduced levels of IL-17A (Fig 8F). Despite normal numbers of RORγt^+^Tbet^+^ CD4^+^ T cells (Fig 8E), IL-17A^+^IFN-γ^+^ cells were significantly reduced in infected ΔIAP1/2 mice (Fig 8F). However, production of IL-22 was not defective in the absence of cIAP1 and cIAP2 (Fig 8F). Collectively, our data suggest that cIAP1 and cIAP2 are required within the ILC3 and T_H_17 compartments to control intestinal bacterial infection.

## Discussion

In the present study we demonstrate that the E3 ubiquitin ligases cIAP1 and cIAP2 are necessary for γδT17 cells to transition through to prepubescent life by regulating cytokine-mediated proliferation and stable expression of the lineage defining transcription factors cMAF and RORγt. Thus, during aging, cIAP1 and cIAP2 are required in a cell-intrinsic manner to maintain cMAF and RORγt levels and to allow cells to enter cell cycle in response to IL-7, IL-1β and IL-23. Consequently, γδT17-driven inflammatory responses in the skin and draining LNs of prepubescent ΔIAP1/2 mice are blunted despite normal cell numbers, while by the time animals reach adulthood, γδT17 populations are deficient in gut, skin and LNs. Mechanistically, our data suggest that this is independent of TNF and TNFR1 and most likely through overt activation of the non-canonical NF-κB pathway. Similar to γδT17, ILC3 required cIAP1 and cIAP2 expression during the post-weaning period in order to expand, be maintained until adult life, and induce formation of intestinal ILFs. The ILC3 deficit in ΔIAP1/2 mice together with a defective T_H_17 response, correlated with a profound inability to control intestinal bacterial infection.

The IL-17-producing γδ T cell subset is an innate-like unconventional lymphocyte that is important in many immunological processes ranging from anti-microbial protection to pathogenic inflammation and cancer (Patil *et al*, 2015). γδT17 cells are pre-programmed and functionally mature in the embryonic thymus in mouse and human (Ribot *et al*, 2009; Haas *et al*, 2012). They are exported into peripheral and secondary lymphoid tissues after birth, and evidence suggests that they go through a second wave of transcriptional and functional programming during neonatal life within the tissues they occupy (Kadekar *et al*, 2020; Wiede *et al*, 2017). The molecular cues that γδT17 cells receive within the tissues during that period are obscure. Our data show that the E3 ligases cIAP1 and cIAP2 are required during late neonatal and early prepubescent life in a cell-intrinsic manner for cytokine-induced proliferation, to sustain transcriptional stability and allow optimal inflammatory responses. This work establishes cIAP1/2 as critical molecular regulators of committed tissue-resident γδT17 cells, and underpins the existence and importance of post-thymic temporal events necessary for these cells to be maintained during aging.

cIAP1/2 are central for TNFR1 induced canonical NF-κB activation and cell death (Mahoney *et al*, 2008) , and necessary to suppress overt non-canonical NF-κB signaling (Vallabhapurapu *et al*, 2008; Zarnegar *et al*, 2008) . TNFR1 induced apoptosis and necroptosis are fundamental biological processes regulating cell growth during development, homeostatic turnover and even inflammatory diseases (Kalliolias & Ivashkiv, 2016). The two NF-κB pathways on the other hand are synonymous with cell survival, proliferation and differentiation in ubiquitous cell populations (Hayden & Ghosh, 2011). In T cells they are mostly active downstream of TNF superfamily receptors and the TCR (Oh & Ghosh, 2013). Although the role of several TNF superfamily receptors and ligands have been studied in γδ T cells (Powolny-Budnicka *et al*, 2011; Shibata *et al*, 2011; Silva-Santos *et al*, 2005), the importance of the signaling components of the NF-κB pathway had not been thoroughly investigated. Genetic and pharmacological perturbations of the TNFR1 signaling, combined with aberrant nuclear translocation of RelB that it is the cIAP1/2-mediated control of non-canonical NF-κB that is required for the maintenance of γδT17 cells. This agrees with CD27, a TNF superfamily receptor and potent activator of non-canonical NF-κB (Ramakrishnan *et al*, 2004), suppressing the γδT17 differentiation program (Ribot *et al*, 2009). Importantly, deletion of NIK, the kinase targeted by cIAP1/2 and responsible for activating the non-canonical NF-κB cascade, did not affect γδT17 cell development or homeostasis (Mair *et al*, 2015). This strongly suggests that it is the “brake” imposed by cIAP1/2 in order to avoid over activation of non-canonical NF-κB that is critical and not its baseline activity.

There are a number of transcription factors that are important for the development of γδT17 cells (Parker & Ciofani, 2020) . Ciofani and co-workers showed that cMAF acts early in embryogenesis to allow robust expression of RORγt and thus promote specification and stability of the γδT17 lineage (Zuberbuehler *et al*, 2019). How cMAF and RORγt expression is regulated, however, in γδT17 cells is not well-defined. Herein, we demonstrate that loss of cIAP1/2 results in the progressive downmodulation of cMAF and RORγt after birth, providing a molecular understanding of how lineage defining transcription factors are regulated in these cells. Although loss of cMAF and RORγt during embryonic development resulted in rapid loss of γδT17 cells or their progenitors in the thymus (Zuberbuehler *et al*, 2019; Shibata *et al*, 2011) , we observed that in ΔIAP1/2 mice, cells persist for at least 2 weeks without either transcription factor. Thus, it appears that during neonatal life the impact of cMAF and RORγt in γδT17 cells is less pronounced. The exact molecular steps leading to cIAP1/2-dependent regulation of cMAF and RORγt are currently unclear. Our previous work showed that following cIAP1/2 inhibition, NIK-mediated RelB nuclear translocation suppressed expression of cMAF in T_H_17 cells (Rizk *et al*, 2019). It is plausible therefore, that accumulation of non-canonical NF-κB signaling directly suppresses cMAF, which subsequently suppresses RORγt. Intriguingly, cytokine stimulation and inflammation could partially restore expression of cMAF and RORγt, indicating a certain degree of transcriptional plasticity.

In addition to γδT17 cells, cIAP1/2 were necessary for the establishment of a normal ILC3 population during the post-weaning period in the gut and the formation of ILFs, as well as protection from intestinal extracellular bacterial infection. During infection, we additionally found that cIAP1/2, were critical for the generation of IL-17^+^ and IL-17^+^IFN-γ^+^ CD4+ T cells, which have been associated with protection against pathogens, or tissue damage in the context of inflammation (Omenetti *et al*, 2019). We and others have previously reported that the cIAP-non-canonical NF-κB axis is necessary for T_H_17 differentiation and successful IL-17-driven responses (Rizk *et al*, 2019; Kawalkowska *et al*, 2019), while NIK was shown to be important for the generation of neuropathogenic T_H_17 cells (Lacher *et al*, 2018). Moreover, NIK expression and activation of the non-canonical NF-κB pathway in dendritic cells indirectly regulates maintenance of both T_H_17 cells and ILC3 (Jie *et al*, 2018). Our current data, extend and broaden the immunological importance of this pathway. We would like to propose that through regulation of non-canonical NF-κB, cIAP1/2 are master regulators of innate and adaptive type-3 immunity. Their requirement is necessary during neonatal life to establish functional innate and innate-like type-3 immune cell populations, whereas in the adult they support differentiation of antigen-dependent adaptive type-3 cells.

## Materials and methods

### Mice

All animals were bred and maintained in-house at DTU health tech with the approval of the Danish animal experiments inspectorate. cIAP1^f/f^ and cIAP1^f/f^ cIAP2^-/-^ mice were provided by Prof. W. Wei-Lynn Wong at the University of Zurich, Switzerland with the permission of Prof. John Silke, VIC Australia. RORγt^CRE^ mice were provided by Prof. Gerard Eberl at Pasteur Institute, Paris, France. ROSA26-floxSTOPflox-RFP mice were from the Swiss Immunological Mouse Repository (SwImMR). Lymph nodes from TNFR1^-/-^ mice were provided by Prof. William Agace at Lund University, Sweden, while Lymph nodes from cIAP1^UBA^ mutant mice were provided by Prof. Pascal Meier at The Institute of Cancer research, UK.

### Cell culture media and buffers

For all preparations of single cell suspensions and cell cultures RPMI 1460 (Invitrogen) supplemented with 10% heat inactivated FBS (GIBCO), 20mM Hepes pH 7.4 (Gibco), 50 µM 2-mercaptoethanol, 2 mM L-glutamine (Gibco) and 10,000 U/ mL penicillin-streptomycin (Gibco), was used. Where indicated, IMDM (Invitrogen) was used instead of RPMI 1460 and supplemented as aforementioned. FACS buffer was prepared by supplementing PBS with 3% heat inactivated FBS.

### Lymphocyte isolation from mouse organs

Lymphocytes were isolated from peripheral lymph nodes (axial, brachial and inguinal), thymus, ear skin, small intestinal and colonic lamina propria following the previously described protocols (Kadekar *et al*, 2020). Lymphocytes were isolated from cervical and auricular lymph nodes in case of IMQ-induced psoriasis.

### Ex-vivo culturing of lymphocytes

For staining of cytokines from lymphocytes that were isolated from peripheral lymph nodes of untreated mice, the cells were plated at a denisty of 10x10^6^ cells /ml in 1ml of supplemented RPMI in 12 well plates. The cells were restimulated with 50ng/ml PMA (phorbol myristate acetate; Sigma Aldrich), 750 ng/ml Ionomycin (Sigma Aldrich) and BD GolgiStop (containing monensin at 1:1000 dilution, BD) and cultured for 3.5 hours at 37°C. For estimation of IL-22 production by CD4^+^ and γδ^+^ T cells from homeostatic mice, the lymphocytes were first cultured onvernight with 40ng/ml rmIL-23 (R&D) the restimulated with PMA, Ionomycing and BD GolgiStop as aforementioned. The cells were then harvested and used for flow cytometry staining.

In case of lymphocytes that were isolated from peripheral lymph nodes or skin in IMQ-experiments, the cells were plated at a denisty of 5x10^6^ cells /ml in 1ml of supplemented IMDM in 24 well plates. The cells were restimulated with 50ng/ml PMA (phorbol myristate acetate; Sigma Aldrich), 750 ng/ml Ionomycin (Sigma Aldrich) and BD Golgiplug (containing Brefeldin A at 1:1000 dilution, BD) and cultured for 3.5 hours at 37°C. The cells were then harvested and used for flow cytometry staining.

Alternatively, lymphocytes that were isolated from mesenteric lymph nodes or colonic lamina propria in *Citrobacter rodentium* infection experiments, the cells were plated at a denisty of 5x10^6^ cells /ml in 1ml of supplemented IMDM in 24 well plates. The cells were subsequently treated with 40ng/ml rmIL-23 (R&D) for 3 hours, followed by 50ng/ml PMA (phorbol myristate acetate; Sigma Aldrich), 750 ng/ml Ionomycin (Sigma Aldrich) and BD Golgiplug (containing Brefeldin A at 1:1000 dilution, BD) and cultured for an additional 3.5 hours at 37°C.

For cell cycle assay experiments, lymphocytes that were isolated from peripheral lymph nodes of mice, were plated at a denisty of 5x10^6^ cells /ml in 1ml of supplemented RMPI in 24 well plates. The cells were treated with either 20ng/ml rmIL-7 (R&D), or with 10 ng/ml rmIL-1β (Biolegend) + 20 ng/ml rmIL-23 (R&D), or 20 ng/ml rhIL-2 (Biolegend), or anti-CD3 (2 μg/ml; clone 145-2C11) for 48 hours. The cells were subsequently harvested for flow cytometry staining.

### IMQ-induced psoriasis

Psoriasis was induced in mice by applying 7 mg of Aldara cream (containing 5% imiquimod) to the dorsal side of each ear for 7 days. Histological sections were prepared by fixing ear tissue in 10% formalin overnight and then paraffin embedded. The paraffin embedded sections were cut and stained by H&E.

### Flow cytometry staining

Surface antigens, intracellular cytokines and cell cycle assay were stained for flow cytometry as previously described (Rizk *et al*, 2019). For transcription factor staining, the cells were first stained for live/dead discrimination followed by surface antigen staining and subsequently fixed using Foxp3 fixation/permeabilization buffer (Thermo Fisher) for 1 hour at 4°C. The cells were washed once with then stained with the desired antibodies in Foxp3 perm/wash buffer for 1 hour at 4°C. The cells were washed once again and resuspended in FACS buffer and analyzed using BD LSRFortessa. The following antibodies were used herein at 1:200 dilution unless otherwise indicated: Fixed viability stain-700 (FVS700, BD, 1:1000), anti-IL-17A (TC11-18H10; BV786 and PE), anti-IFNγ (XMG1.2; PE-Cy7, APC, BV711 and Percp-cy5.5), anti-IL-22 (1H8PWSR; PE), anti-cMAF (symOF1; PE, eF660 or Percp-Cy5.5; 5 μL/test), anti-CD4 (GK1.5; BUV395 and FITC), anti-TCRγδ (GL3; BV421 and APC), anti-KLRG1 ( 2F1 , BV786), anti-CD27 (LG.3A10; PE-Cy7 and BV650), anti-CCR6 (140706; Alexa Fluor 647), anti-CD44 (1M7; V500), anti-CD19 (6D5; FITC), anti-TCRβ (H57-597; APC-eflour780), anti-CD3e (145-2C11, PeCF594 and PE), anti-Tbet(4B10; PeCy7), anti-CD8 (53-6.7;FITC), anti-Vγ5 (536; FITC), anti-Vγ4(UC3-10A6; Percp-eflour710), anti-GATA3(TWAJ; Percp-eFlour710; 1:30), anti-CD45(30-F11;PE and V500), anti-CD127 (SB/199; BUV737) and anti-RORγt (B2D; APC and PE).

### Administration of Anti-TNF

For neutralization of TNF, 1 week old pups were weighed and i.p. injected with the a-TNF (Adalimumab, brand name HUMIRA) at 5 mg/kg body weight once a week until weaning. After weaning the mice were i.p. injected with 10 mg/kg body weight twice a week until euthanasia at approximately 12 weeks of age.

### Transfer of neonatal γδ T cells to RAG1^-/-^ hosts

First, thymi from 1-2 days old mice were isolated and crushed individually against 70 μm filter to prepare single cell solutions. Subsequently, total γδ T cells were enriched by magnetic depletion of CD4^+^, CD8^+^, TCRβ^+^ cells as follows: total thymocytes were re-suspended in MACS buffer at 1e8 cells/ml containing 50 μL/ml normal rat serum and 1:200 biotin labelled anti-CD4^+^ (GK1.5), CD8^+^ (53-6.7) and TCRβ^+^ (H57-597) antibodies; the cells were incubated for 10 minutes at room temperature and then incubated with 75 μL/ml EasySep RaphidSphere streptavidin beads (#50001) for 2.5 minutes then transferred to EasySep magnet for 2.5 minutes. The non-bound fraction was collected by decantation and centrifugated for 5 minutes at 400g at 4°C.. The enriched γδ T cells from each donor mouse were re-suspended in PBS and then i.v. injected into the tail vein of a RAG1^-/-^ host. The RAG1^-/-^ hosts were euthanized for collection of organs after 12 weeks.

### Bone marrow chimeras

The bone marrow cells for reconstitution were isolated by flushing the tibia and femur, which were dissected from donor mice, with culture media. Total bone marrow cells were then centrifuged at 400g for 5 minutes at 4°C. The cells were then re-suspended and passed through 70 μm filter. Subsequently, red blood cells were then lysed using RBC lysis buffer (Biolegend) and a single cell suspension of bone marrow cells was the prepared by passing the cells through 40 μm filter. The prepared cells were then counted and mixed as appropriate.

Conversely, host mice were sub-lethally irradiated by 2 doses of 4.5 Gy that were at least 4 hours apart. After 24 hours, the hosts were reconstituted with 10e6 bone marrow cells that were i.v. injected into the tail vein of the host mice. The hosts were euthanized for organs after at least 12 weeks.

### Immunofluorescent imaging of intestinal tissue

To assess the presence of SILT in the intestines of WT or ΔIAP1/2 by confocal laser microscopy, the distal ileum was taken and flushed once with HBSS (Thermo Fisher) to remove intestinal contents. Cleaned intestines were fixed for 8h in 4% PFA (Sigma-Aldrich) in PBS and stored in washing buffer (PBS+5%FCS+0.2% Triton X-100 (Sigma-Aldrich)+0.01% Thimerosal (Sigma-Aldrich)) until further use. To prepare the collected intestines for staining, tissues were embedded in 4% UltraPure™ Low Melting Point Agarose (Thermo Fisher) in PBS, sectioned with a swinging blade microtome (Leica VT1200S) into 50 micron sections and permeabilized overnight using the Foxp3 Transcription Factor Staining Buffer Set (Thermo Fisher). Permeabilized sections were stained in the supplied permbuffer with an antibody against RORγt (AFKJS-9; unconjugated), followed by a washing step in permbuffer and incubation with a biotinylated secondary antibody against the primary anti-RORγt antibody (Biotinylated anti-rat; Jackson ImmunoResearch). To detect RORγt^+^ ILC and B cells, sections were washed again in permbuffer and incubated in permbuffer with antibody against B220 (RA3-6B2; AF647) and streptavidin-conjugated AF555 (Thermo Fisher), as well as DAPI (Thermo Fisher) to stain all nucleated cells. Sections were washed one more time, mounted on glass slides with ProLong Gold (Thermo Fisher) and analyzed using an LSM710 confocal laser microscope (Carl Zeiss). Images of ≥5 different sections per mouse were acquired with the Zeiss Zen v2.3 software (Carl Zeiss) and analyzed using Imaris v8 (Bitplane/Oxford Instruments) and Fiji v2.1.0/1.53c (Schindelin *et al*, 2012).

### Murine Citrobacter rodentium infection

Starter cultures of *Citrobacter rodentium* strain DBS100 (ATCC 51459; American Type Culture Collection) were grown overnight at 37°C in Luria-Bertani (LB) medium. The cultures were then used at 5% v/v to inoculate sterile LB medium. The cultures were grown at at 37°C to an OD600 of 0.8-1 and the CFU count was determined from the OD600 measurement using the following formula: CFU/ml =(5x10^8^)(OD)–3x10^7^. Subsequently, the bacteria was collected by centrifugation at 4000g for 10 minutes. The bacterial pellet was then resuspended in LB medium to give at 2x10^9^ CFU/100 μL. To infect adult mice, the mice were orally gavaged with either 100 μL of *Citrobacter rodentium* or LB control. The mice were weighed before oral gavage and once daily until termination of the experiment. At day 12 post infection all mice were euthanized, dissected to collect organs and fecal samples.

The collected fecal samples were weighed and dissolved in PBS and then serially diluted. The serial dilutions were plated on Brilliance™ E. coli/coliform Agar (CM0956, Thermo Fisher) and incubated overnight at 37°C. *Citrobacter rodentium* colonies were identified as being pink colonies and enumerated, while E. coli colonies were identified as purple colonies. CFU/g stool was then calculated as previously described (Bouladoux *et al*, 2017).

### Immunofluorescent imaging of nuclear RelB in γδ T cells

Total lymphocytes were isolated from peripheral (axial, brachial and inguinal), cervical and auricular lymph nodes of 4 weeks old mice as described above. Subsequently, total γδ T cells were enriched by magnetic depletion of CD4^+^, CD8^+^, CD19^+^ and TCRβ^+^ cells as aforementioned. The cells were then stained with FVS700 for discrimination of live and dead cells and then stained for surface antigens with the following antibodies: anti-TCRγδ (GL3; APC), anti-CD27 (LG.3A10; PE-Cy7) and anti-TCRβ (H57-597; APC-eflour780) all at 1:200 dilution. The cells were subsequently sorted into TCRγδ^+^ CD27^+^ or TCRγδ^+^ CD27^-^ cells using BD ARIA-FUSION cell sorter. The sorted cells were collected into cell culture medium and centrifuged at 400g for 5 minutes at 4°C. The cells were then fixed using Foxp3 fixation/permeabilization buffer (Thermo Fisher) for 1 hour at 4°C. The cells were washed once with then stained with Foxp3 perm/wash buffer containing anti-TCRγδ (GL3; APC, 1:50), anti-CD3e (145-2C11 or 17A2; biotin, 1:100) and anti-RelB (D-4, Santa-cruz, 1:40) for 1-hour 4°C in Foxp3 perm/wash buffer. Again, the cells were washed once and stained with streptavidin-conjugated AF488 (Biolegend, 1:100) and anti-mouse AF555 (1:100) for 1 hour 4°C in Foxp3 perm/wash buffer. The cells were then washed once more as previous and stained with DAPI to highlight cell nuclei and washed once more with PBS. Washed cells were mounted on a glass slide using ProLong Gold and imaged with an LSM710 confocal laser microscope and were acquired and analyzed with the Zen v2.3 software and and Fiji v2.1.0/1.53c (Schindelin *et al*, 2012).

### Real-Time quantitative PCR

At the indicated timepoints, γδ T cells were sorted from the thymus or lymph nodes of mice directly into RLT buffer mixed with β-mercaptoethanol. RNA was then extracted from using Qiagen RNAeasy microkit following the manufacturer’s instructions.

Subsequently, cDNA was prepared using BioRad Iscript cDNA synthesis kit using the manufacturer’s protocol. Gene expression was then measured by RT-qPCR reactions using BioRad SSOFast EvaGreen supermix, which were run on CFX96 (Biorad) and analyzed using Bio-rad CFX manager software. The following primers were used for RT-qPCR: Actb, Fwd-GGCTGTATTCCCCTCCATCG, Rev-CCAGTTGGTAACAATGCCATGT; RelB, Fwd-GCTGGGAATTGACCCCTACA, Rev-CATGTCGACCTCCTGATGGTT; Birc2, Fwd-TGCCTGTGGTGGGAAACTGA, Rev-GCTCGGGTGAACAGGAACA.

