## Supplementary figures for "The cIAP ubiquitin ligases sustain type 3 γδ T and innate lymphoid cells during aging to allow normal cutaneous and mucosal responses"

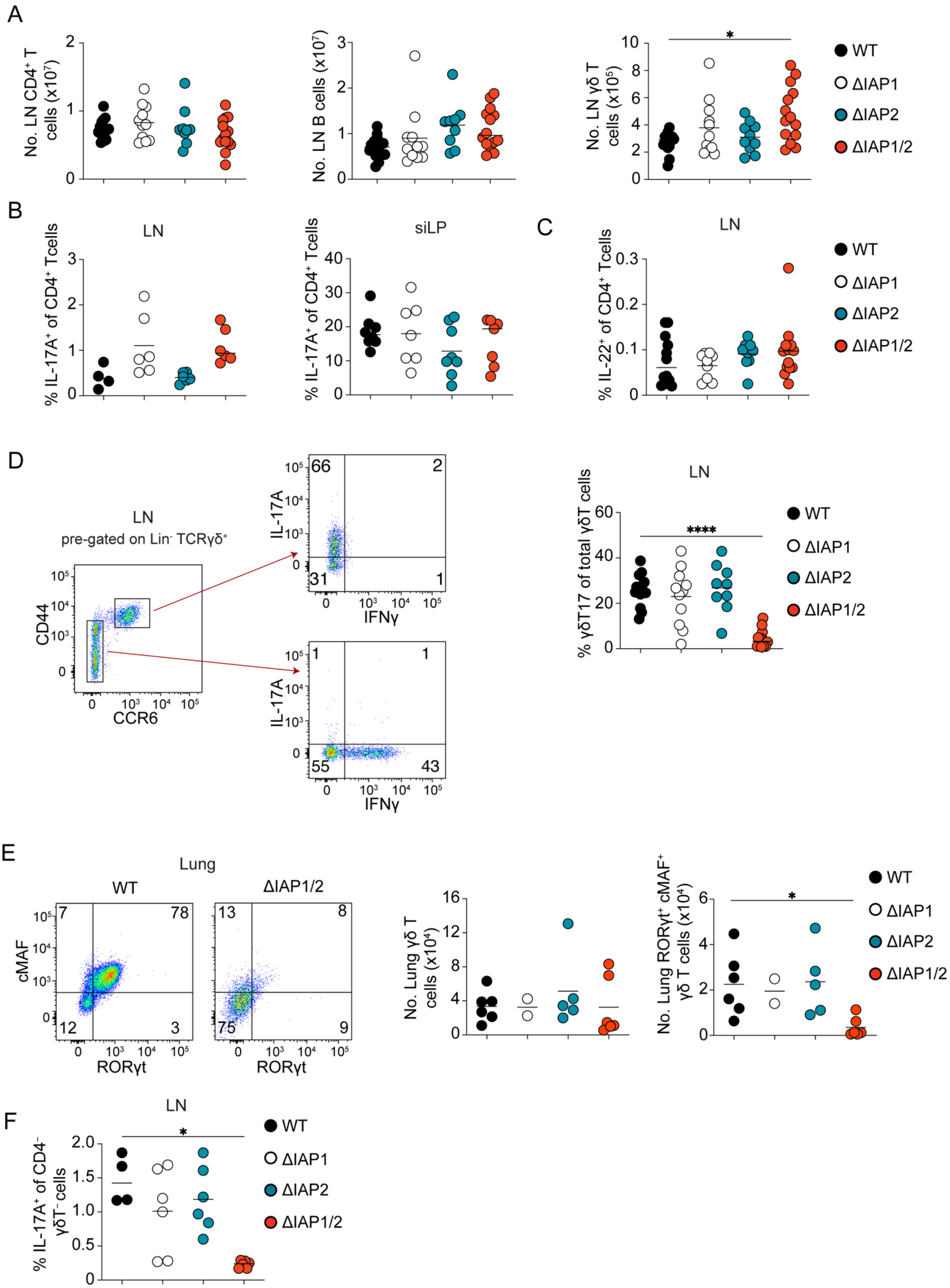
A
No. LN CD4⁺ T cells (x10⁷)
No. LN B cells (x10⁷)
No. LN γδ T cells (x10⁵)
*
WT
ΔIAP1
ΔIAP2
ΔIAP1/2
B
LN
% IL-17A⁺ of CD4⁺ Tcells
siLP
% IL-17A⁺ of CD4⁺ Tcells
C
LN
% IL-22⁺ of CD4⁺ Tcells
D
LN
pre-gated on Lin⁻ TCRγδ⁺
CD44
CCR6
IL-17A
IFNγ
66
2
31
1
1
1
55
43
LN
% γδT17 of total γδT cells
****
E
Lung
WT
ΔIAP1/2
cMAF
RORγt
7
78
12
3
13
8
75
9
No. Lung γδ T cells (x10⁴)
No. Lung RORγt⁺ cMAF⁺ γδ T cells (x10⁴)
*
F
LN
% IL-17A⁺ of CD4⁻ γδT⁻cells
*

**Supplementary Figure 1. cIAP1 and cIAP2 are required for the homeostasis of γδT17 but not B or CD4$^+$ T cell populations in ΔIAP1/2 mice**

(A) Numbers of CD4$^+$ T cells, B cells or total γδ T cells in the LNs of adult WT, ΔIAP1, ΔIAP2 and ΔIAP1/2 mice. In graphs, each symbol represents a mouse, and lines represent the mean, data is pool of 5 experiments. (B) frequency of Il-17A$^+$ cells within CD4$^+$ T cells in the LNs or siLP of adult WT, ΔIAP1, ΔIAP2 and ΔIAP1/2 mice. In graphs, each symbol represents a mouse, and lines represent the mean, data is pool of 3 experiments for LNs analysis and 5 experiments for the siLP analysis. (C) frequency of IL-22$^+$ cells within CD4$^+$ T cells in the LNs of adult WT, ΔIAP1, ΔIAP2 and ΔIAP1/2 mice. In graph, each symbol represents a mouse, and the line represents the mean, data is pool of 4 experiments. (D) Representative flow cytometric analysis of IL-17A and IFNγ production by the subsets of γδ T cells and frequency of γδT17 cells within total γδ T cells in the LNs of adult WT, ΔIAP1, ΔIAP2 and ΔIAP1/2 mice. In graph, each symbol represents a mouse, and the line represents the mean, data is pool of 5 experiments. (E) Numbers of total γδ T and RORγt+ γδ T cells in the lungs of adult WT, ΔIAP1, ΔIAP2 and ΔIAP1/2 mice. In graph, each symbol represents a mouse, and the line represents the mean, , data is pool of 2 experiments. (F) frequency of Il-17A$^+$ cells within CD4$^-$ T cells in the LNs of adult WT, ΔIAP1, ΔIAP2 and ΔIAP1/2 mice. In graphs, each symbol represents a mouse, and lines represent the mean, data is pool of 3 experiments for LNs analysis. *$P < 0.05$, ****$P < 0.0001$ using Kruskal-Wallis test with Dunn's correction.

A

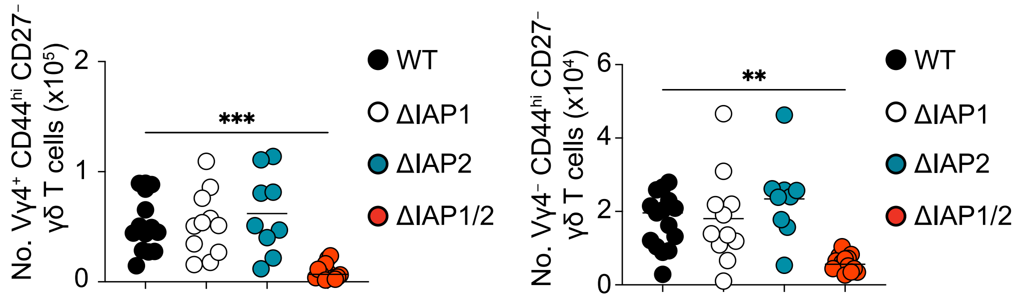


**Supplementary Figure 2. cIAP1 and cIAP2 are redundantly required for maintenance of γδT17 cell subsets in the LNs.**

(A) numbers of Vγ4$^{+}$ and Vγ4$^{-}$ γδT17 cells in the LNs of WT, ΔIAP1, ΔIAP2 and ΔIAP1/2 mice. In graphs, each symbol represents a mouse, and lines represent the mean, data is pool of 5 experiments. **P < 0.01, ***P < 0.001 using Kruskal-Wallis test with Dunn's correction.

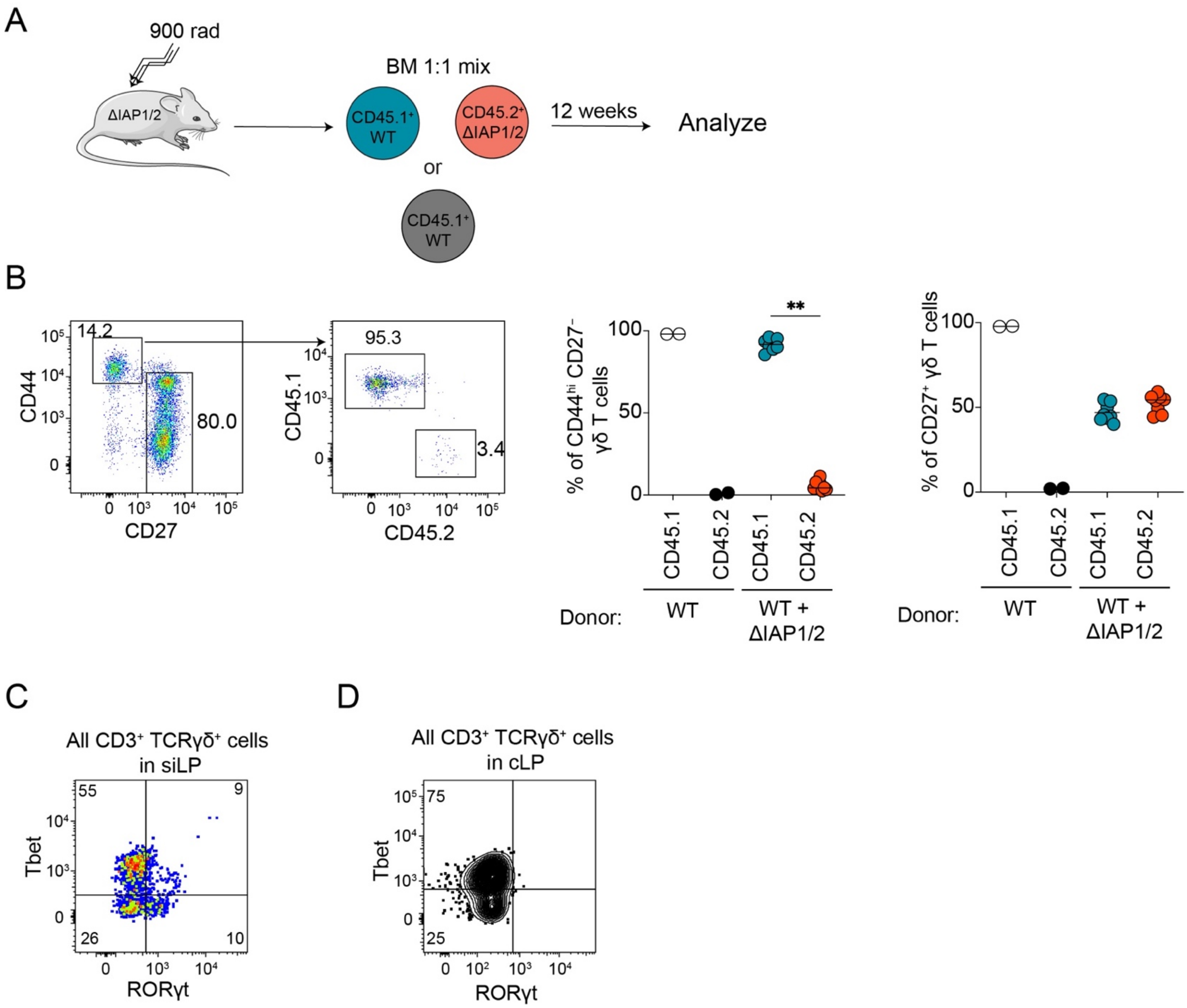


**Supplementary Figure 3. cIAP1 and cIAP2 are intrinsically required for the homeostasis of γδT17 cells.**

(A) Graphical representation of the experimental setup for bone marrow chimera experiments using ΔIAP2 mice as hosts. (B) representative flow cytometric analysis (dot plots) and frequency (graphs) of WT ($CD45.1^+$) or ΔIAP1/2 ($CD45.2^+$) -derived γδT17 and $CD27^+$ γδ T cells within γδ T cells population in the LNs of bone marrow reconstituted ΔIAP1/2 hosts. In graphs, each symbol represents a mouse, and lines represent the mean, data is pool of 2 experiments. **P < 0.01 using Wilcoxon-rank t-test. (C) representative flow cytometric analysis of RORγt$^+$ Tbet$^+$ γδ T cells in the siLP of host ΔIAP1/2 (CD45.2$^+$) mice follwing bone marrow reconstitution. (D) representative Flow cytometric analysis RORγt$^+$ Tbet$^+$ γδ T cells in the cLP of RAG1$^{-/-}$ hosts after transfer of neonatal γδ T cells from WT pups.

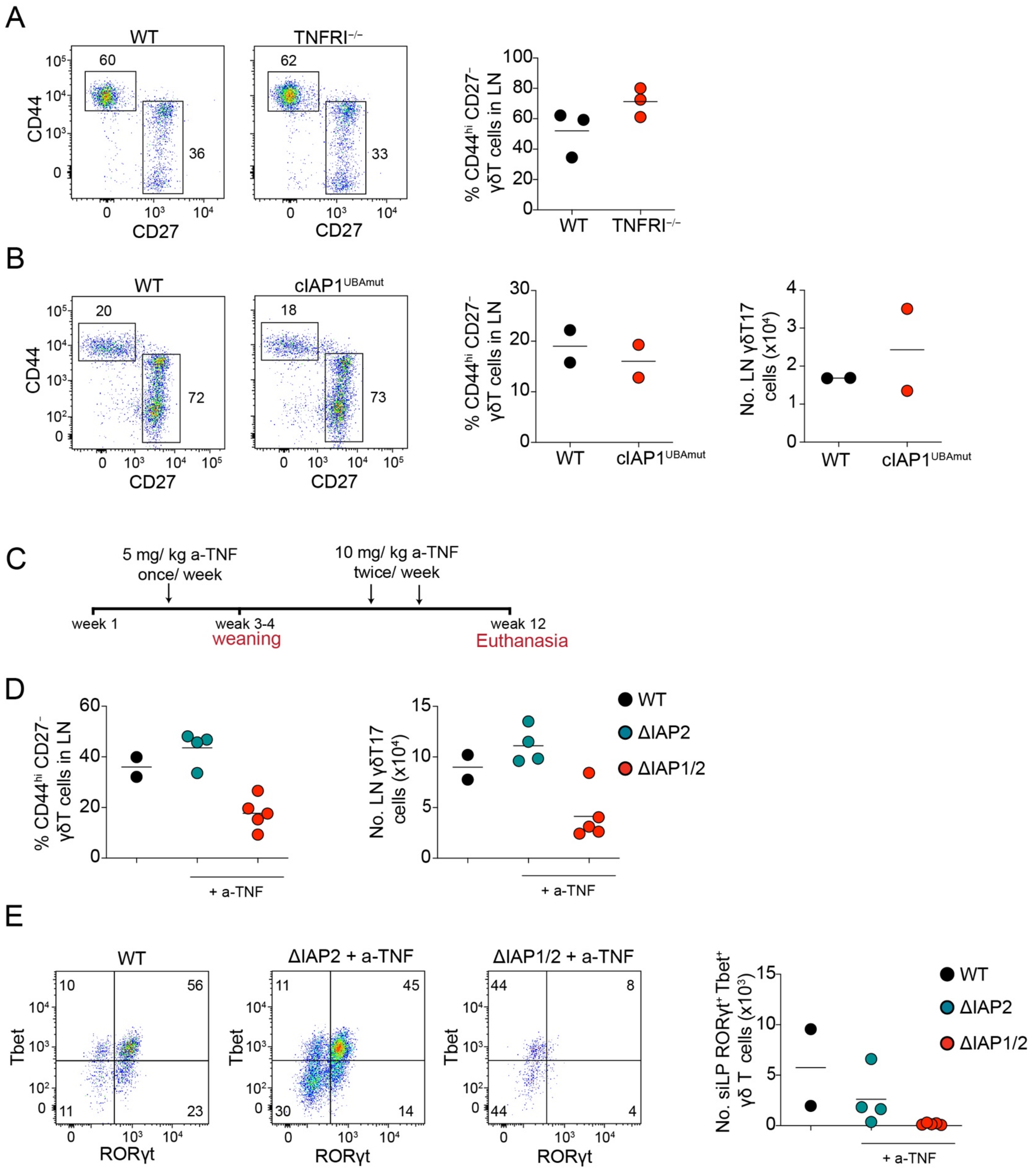


**Supplementary Figure 4. The defect in γδT17 cells from ΔIAP1/2 mice is independent of TNFRI induced cell death.**

(A) representative flow cytometric analysis (dot plots) and frequency (graphs) of γδT17 cells in the LNs of TNFRI$^{-/-}$ mice or littermate controls. (B) representative flow cytometric analysis (dot plots), frequency and numbers (graphs) of γδT17 cells in the LNs of cIAP1$^{UBAmut}$ mice or littermate controls. For graphs in (A-B), each symbol represents a mouse, and lines represent the mean. (C) Graphical representation of the experimental setup for TNF-neutralization experiments. (D) frequency and numbers of γδT17 cells in the LNs of a-TNF-treated ΔIAP1/2 and ΔIAP2 mice or WT control mice. In graphs, each symbol represents a mouse, and lines represent the mean, data is pool of 2 experiments. (E) representative flow cytometric analysis (dot plots) and numbers (graph) of RORγt$^{+}$ Tbet$^{+}$ γδ T cells in the siLP of a-TNF-treated ΔIAP1/2 and ΔIAP2 mice or WT control mice. In graph, each symbol represents a mouse, and lines represent the mean, data is pool of 2 experiments.

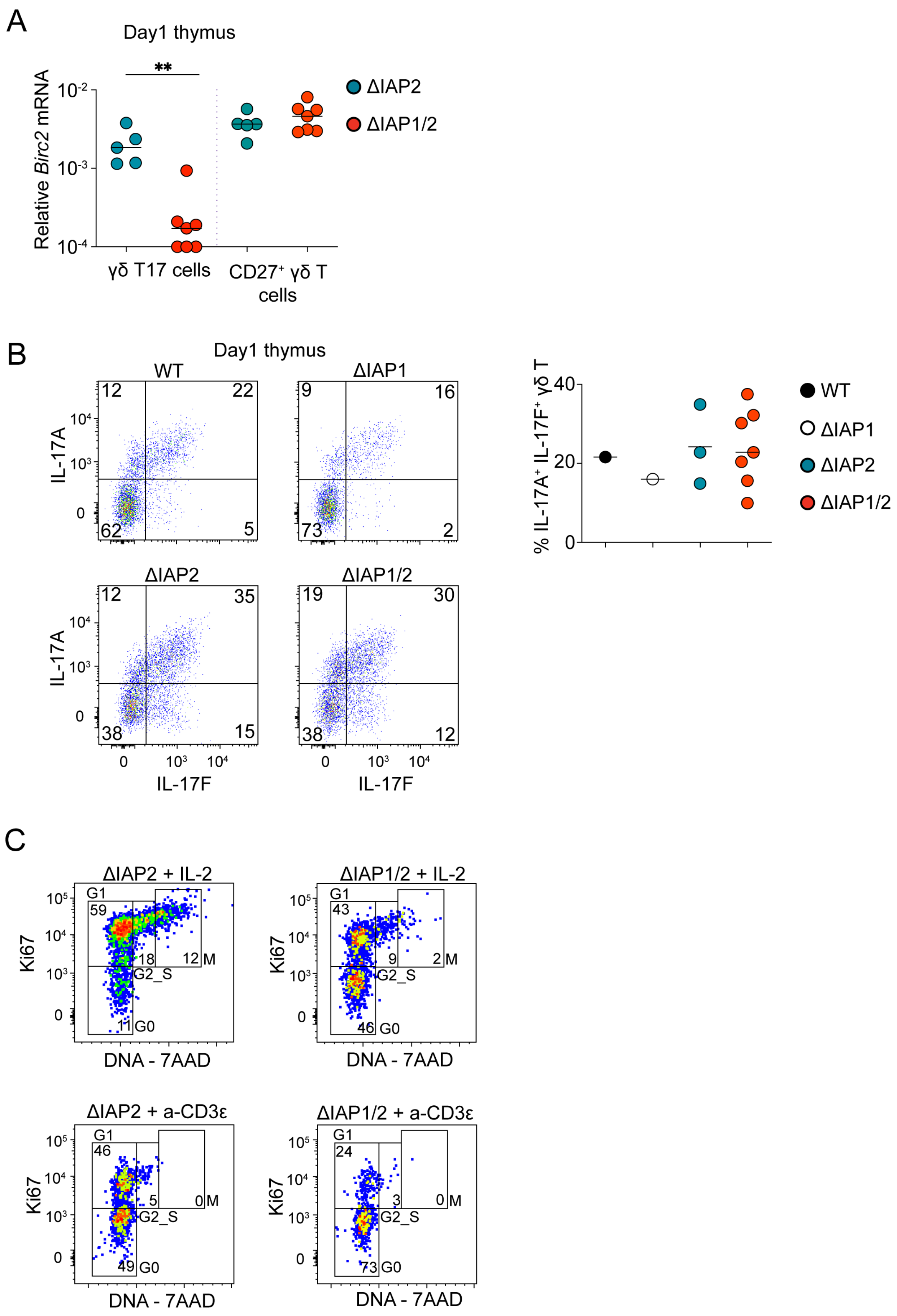
A
Day1 thymus
**
Relative Birc2 mRNA
10-2
10-3
10-4
γδ T17 cells
CD27+ γδ T cells
ΔIAP2
ΔIAP1/2
B
Day1 thymus
WT
ΔIAP1
ΔIAP2
ΔIAP1/2
IL-17A
IL-17F
12 22 62 5
9 16 73 2
12 35 38 15
19 30 38 12
% IL-17A+ IL-17F+ γδ T
WT
ΔIAP1
ΔIAP2
ΔIAP1/2
C
ΔIAP2 + IL-2
ΔIAP1/2 + IL-2
ΔIAP2 + a-CD3ε
ΔIAP1/2 + a-CD3ε
Ki67
DNA - 7AAD
G1
G2_S
M
G0
59 18 12 11
43 9 2 46
46 5 0 49
24 3 0 73

**Supplementary Figure 5. cIAP1 and cIAP2 are not required for the thymic development of γδT17 cells.**

(A) Quantitative real-time PCR (qRT-PCR) analysis of *Birc2* mRNA in thymic γδT17 and CD27+ γδ T cells of 1-day old ΔIAP2 and ΔIAP1/2 pups. **P < 0.01 using Mann-Whitney t-test. (B) flow cytometric analysis (dot plots) and frequency (graph) of IL-17A$^+$ IL17F$^+$ cells within γδT17 cells in the thymus of 1-day old WT, ΔIAP1, ΔIAP2 and ΔIAP1/2 pups. In graph, each symbol represents a mouse, and lines represent the mean. (C) Representative flow cytometric analysis of cells in G0, G1, S or G2/M cell cycle stages within γδT17 cells that were ex-vivo cultured with either IL-2 or a-CD3ε for 48 hours.

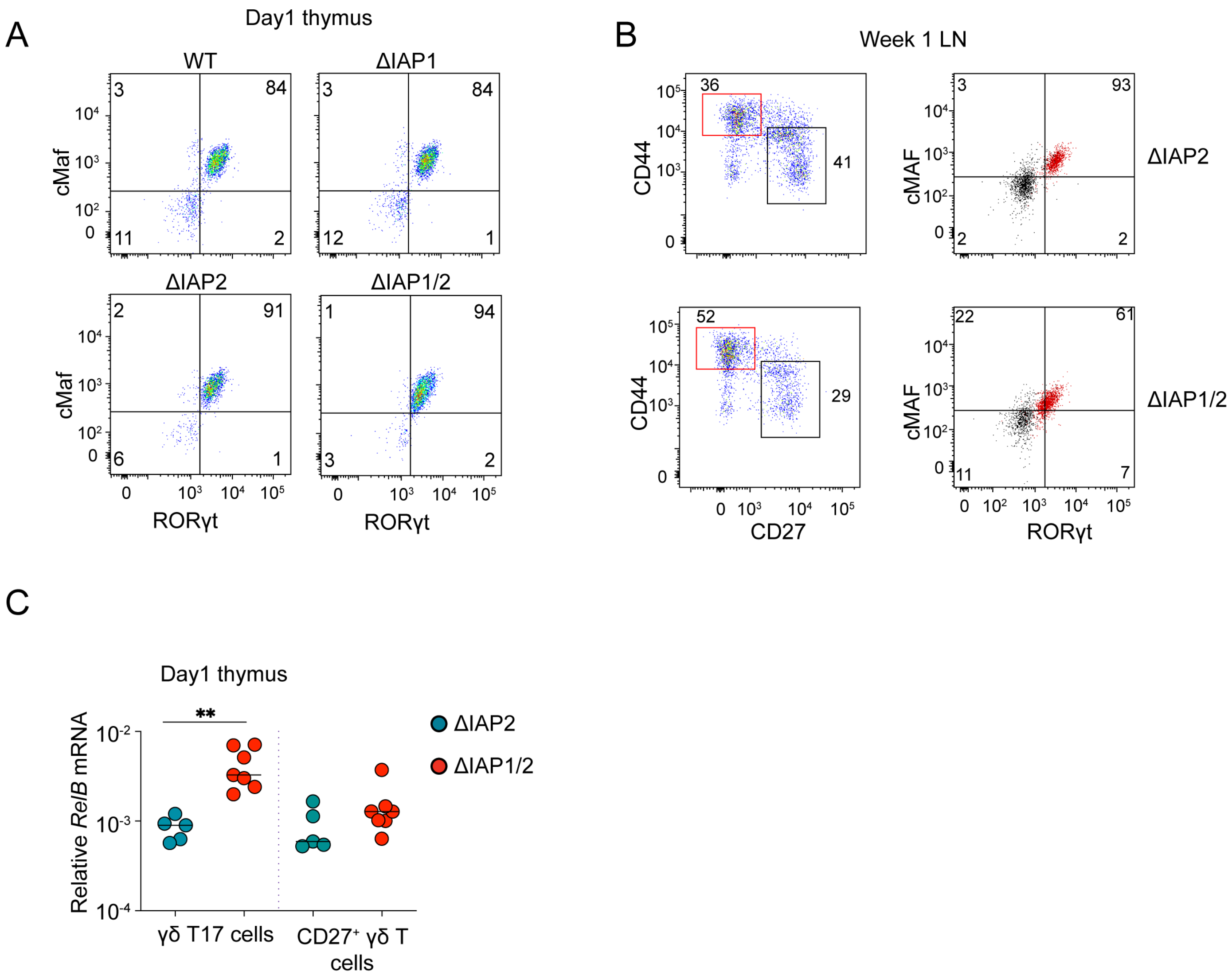


**Supplementary Figure 6. Deficiency of cIAP1 and cIAP2 or RelB upregulation do not affect the thymic development of γδT17 cells.**

(A) flow cytometric analysis of RORγt and cMAF expression by γδT17 in the thymus of 1-day old WT, ΔIAP1, ΔIAP2 and ΔIAP1/2 pups**.** (B) flow cytometric analysis γδ T cell subsets and the expression of RORγt$^{+}$ cMAF$^{+}$ cells by γδT17 cell compared to CD27$^{+}$ γδ T cells in the LNs of 1-week-old ΔIAP2 and ΔIAP1/2 mice. (C) Quantitative real-time PCR (qRT-PCR) analysis of *RelB* mRNA in thymic γδT17 and CD27+ γδ T cells of 1-day old ΔIAP2 and ΔIAP1/2 pups. **P < 0.01 using Mann-Whitney t-test.

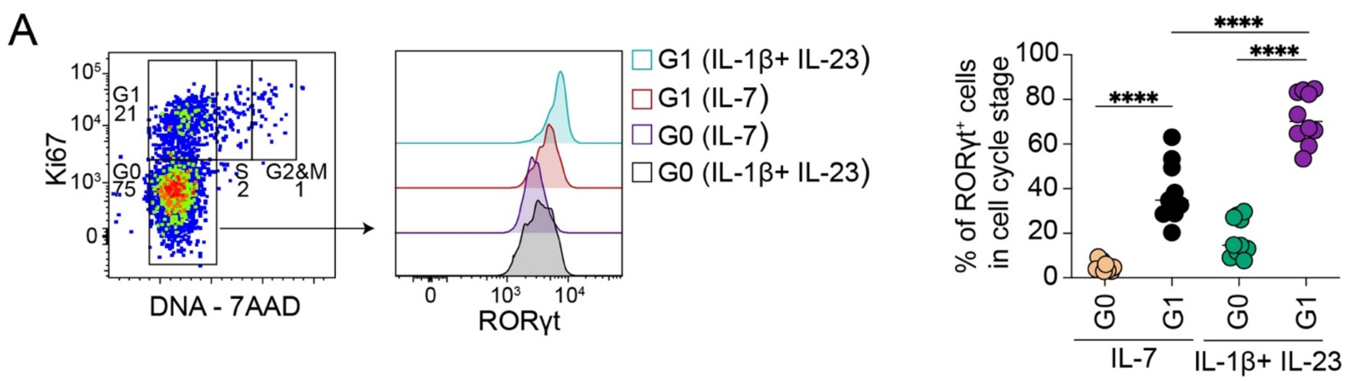


**Supplementary Figure 7. cIAP1/2 deficient γδT17 cells upregulate RORγt upon cell cycle entry.**

(A) Representative flow cytometric analysis (dot plot and histogram) and frequency (graph) of RORγt$^{+}$ in γδT17 cells in G0 or G1 cell cycle stages following ex-vivo culture of lymphocytes from the LNs of 4-week-old ΔIAP1/2 with the indicated cytokines for 48 hours. In graphs, each symbol represents a mouse, and lines represent the mean, data is pool of 3 experiments. ****P < 0.0001 using Mann-Whitney test

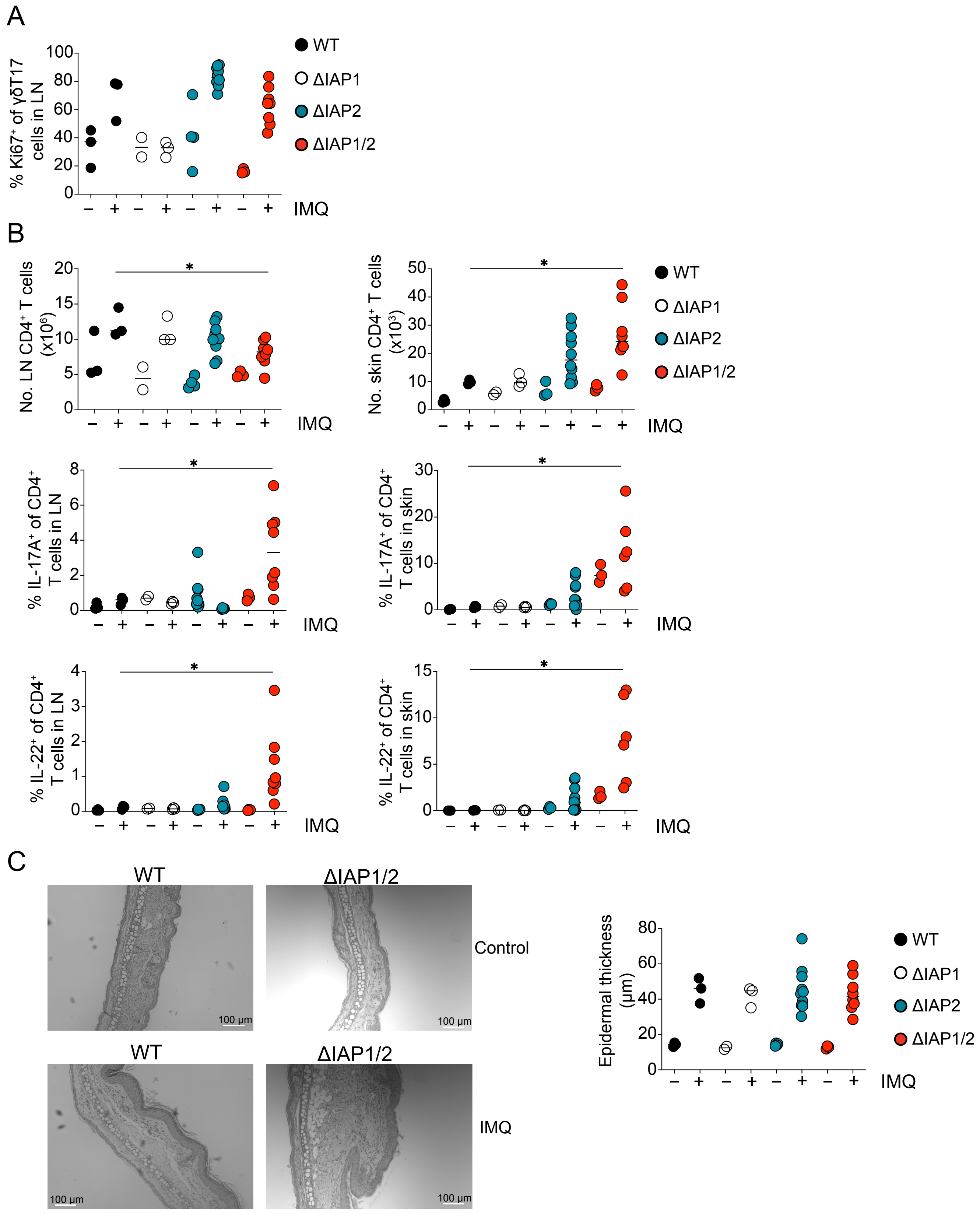


**Supplementary Figure 8. ΔIAP2 mice are not protect from IMQ-induced psoriasis due to intact CD4⁺ T cell response and partial rescue of γδT17 cells.**

(A) Frequency of Ki-67$^+$ cells within γδT17 cells in the LNs of 4-week-old control or IMQ-treated WT, ΔIAP1, ΔIAP2 or ΔIAP1/2 mice. (B) Numbers of CD4$^+$ T cells in the LNs and skin (top graphs), frequencies of IL-17$^+$ (middle graphs) and IL-22+ (bottom graphs) cells within CD4$^+$ T cells in the LNs and skin of 4-week-old control or IMQ-treated WT, ΔIAP1, ΔIAP2 or ΔIAP1/2 mice. In graphs, each symbol represents a mouse, and the line represent the median, data is pool of 3 experiments. *$P < 0.05$, using Mann-Whitney test. (C) representative images and quantification (graph) of epidermal thickening from H&E-stained skin sections of 4-week-old control or IMQ-treated WT or ΔIAP1/2 mice. In graphs, each symbol represents a mouse, and the line represent the median, data is pool of 3 experiments.

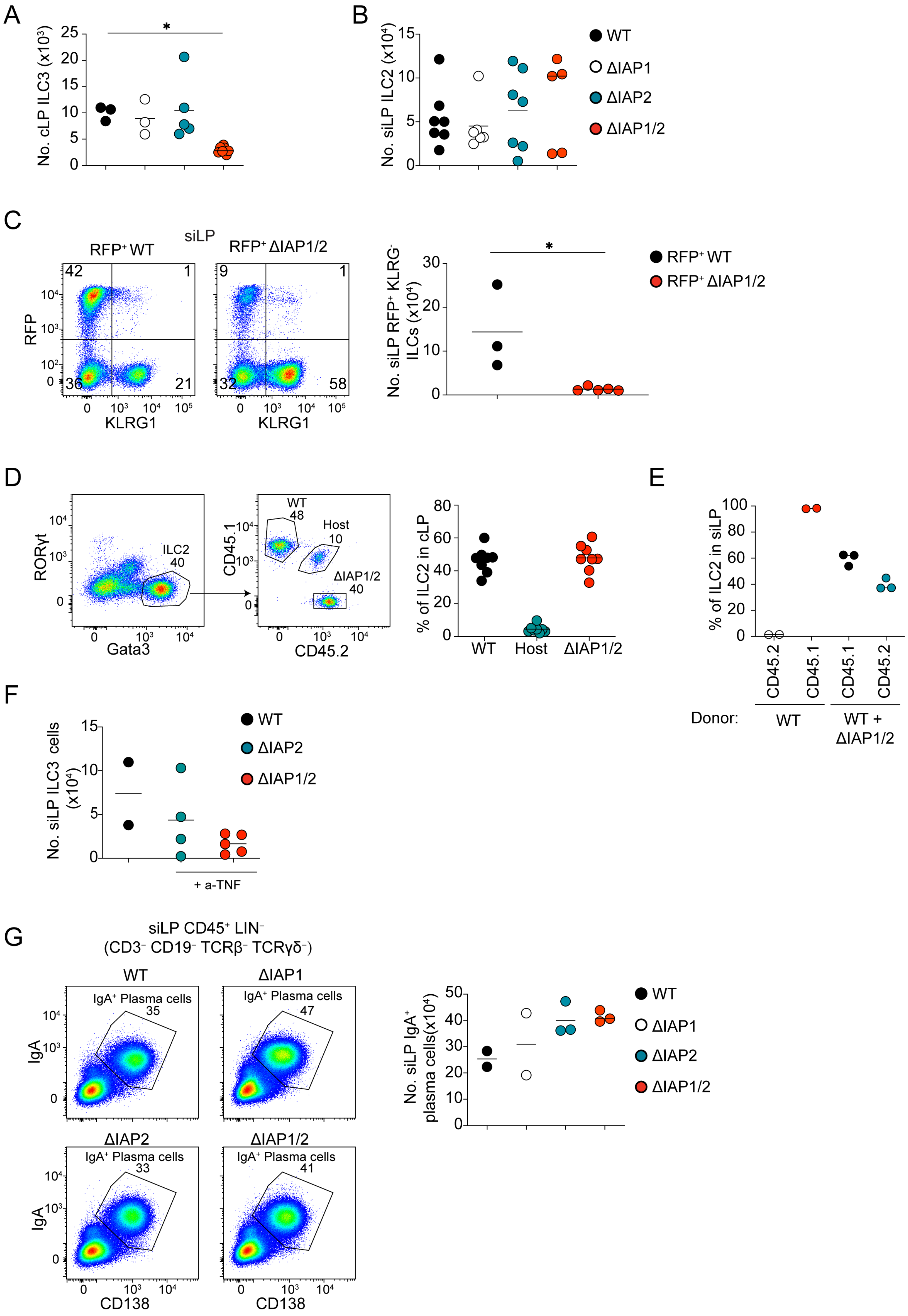
A
No. cLP ILC3 (x10³)
25
20
15
10
5
0
*
B
No. siLP ILC2 (x10⁴)
15
10
5
0
WT
ΔIAP1
ΔIAP2
ΔIAP1/2
C
siLP
RFP⁺ WT
RFP⁺ ΔIAP1/2
RFP
KLRG1
42
1
36
21
9
1
32
58
No. siLP RFP⁺ KLRG⁻ ILCs (x10⁴)
30
20
10
0
*
RFP⁺ WT
RFP⁺ ΔIAP1/2
D
RORγt
Gata3
ILC2
40
CD45.1
CD45.2
WT
48
Host
10
ΔIAP1/2
40
% of ILC2 in cLP
80
60
40
20
0
WT
Host
ΔIAP1/2
E
% of ILC2 in siLP
100
80
60
40
20
0
CD45.2
CD45.1
CD45.1
CD45.2
Donor:
WT
WT + ΔIAP1/2
F
No. siLP ILC3 cells (x10⁴)
15
10
5
0
WT
ΔIAP2
ΔIAP1/2
+ a-TNF
G
siLP CD45⁺ LIN⁻
(CD3⁻ CD19⁻ TCRβ⁻ TCRγδ⁻)
WT
ΔIAP1
ΔIAP2
ΔIAP1/2
IgA⁺ Plasma cells
35
47
33
41
IgA
CD138
No. siLP IgA⁺ plasma cells(x10⁴)
50
40
30
20
10
0
WT
ΔIAP1
ΔIAP2
ΔIAP1/2

**Supplementary Figure 9. Combined cIAP2 deficiency and *Rorc*-driven cIAP1 ablation does not impact intestinal ILC2s or $IgA^+$ plasma cells.**

(A) Numbers of total ILC3s in the cLP of adult WT, ΔIAP1, ΔIAP2 or ΔIAP1/2 mice. In graphs, each symbol represents a mouse, and the line represent the mean, data is pool of 3 experiments. *$P< 0.05$ using Kruskal-Wallis test with Dunn's correction. (B) Numbers of total ILC2s in the siLP of adult WT, ΔIAP1, ΔIAP2 or ΔIAP1/2 mice. In graphs, each symbol represents a mouse, and the line represent the mean, data is pool of 5 experiments. (C) Representative flow cytometry analysis and numbers of $KLRG1^-$ $RFP^+$ ILC in the siLP of RORc-cre x ROSA26-LSL-RFP or ΔIAP1/2 x ROSA26-LSL-RFP mice. *$P< 0.05$ using Mann-Whitney T-test. (D) representative flow cytometric analysis (dot plots) and frequency (graph) of WT ($CD45.1^+$), host ($CD45.1^+$ C $D45.2^+$) or ΔIAP1/2 ($CD45.2^+$)- derived ILC2 in the cLP of bone marrow reconstituted hosts. In graph, each symbol represents a mouse, and lines represent the mean, data is pool of 3 experiments. (E) Frequency of WT ($CD45.1^+$) or ΔIAP1/2($CD45.2^+$) -derived ILC2 in the siLP of bone marrow reconstituted ΔIAP1/2 hosts. In graph, each symbol represents a mouse, and lines represent the mean, data is pool of 2 experiments. (F) numbers of ILC3s in the siLP of a-TNF-treated ΔIAP1/2, ΔIAP2 mice or WT control mice. In graph, each symbol represents a mouse, and lines represent the mean, data is pool of 2 experiments. (G) flow cytometric analysis (dot plots) and numbers (graph) of $IgA^+$ $CD138^+$ plasma cells in the siLP of adult WT, ΔIAP1, ΔIAP2 or ΔIAP1/2 mice. In graph, each symbol represents a mouse, and lines represent the mean, data is pool of 2 experiments.
